# Polo-like kinase 4 homodimerization is not required for catalytic activation, autodestruction, or centriole assembly

**DOI:** 10.1101/2021.11.15.468682

**Authors:** John M. Ryniawec, Daniel W. Buster, Lauren K. Slevin, Cody J. Boese, Anastasia Amoiroglou, Spencer M. Dean, Kevin C. Slep, Gregory C. Rogers

## Abstract

Polo-like kinase 4 (Plk4) is the master-regulator of centriole assembly and cell cycle-dependent regulation of its activity maintains proper centrosome number. During most of the cell cycle, Plk4 levels are nearly undetectable due to its ability to autophosphorylate and trigger its own ubiquitin-mediated degradation. However, during mitotic exit, Plk4 forms a single aggregate on the centriole surface to stimulate centriole duplication. Whereas most Polo-like kinase family members are monomeric, Plk4 is unique because it forms homodimers. Notably, Plk4 *trans*-autophosphorylates a degron near its kinase domain, a critical step in autodestruction. While it is thought that the purpose of homodimerization is to promote *trans*-autophosphorylation, this has not been tested. Here, we generate separation-of-function Plk4 mutants that fail to dimerize, and we show that homodimerization is required to create a binding site for the Plk4 activator, Asterless. Surprisingly, Plk4 dimer mutants are catalytically active in cells, promote centriole assembly, and can *trans*-autophosphorylate by a process based its on concentration-dependent aggregation. Our findings implicate a concentration-dependent pathway of Plk4 activation that does not require Asterless binding or homodimerization. Lastly, we propose a model of Plk4 cell cycle regulation that utilizes both activation pathways – Asterless-dependent and aggregation-driven – to restrict centriole assembly to mother centrioles.

## Introduction

The centrosome is a nonmembrane-bound organelle that functions as the primary microtubule-organizing center in cells and is used to build a variety of microtubule-based protein machines, including the interphase array, mitotic spindles, and cilia (Werner et al., 2017; Kumar and Reiter, 2021). At the centrosome core lies a pair of centrioles, barrel-shaped structures that act as the duplicating elements of the organelle. One centriole is older (the ‘mother’) than the other, having given birth to the younger ‘daughter’ centriole in the previous cell cycle (Nigg and Holland, 2018). Similar to DNA, centrioles duplicate during S-phase when each centriole assembles a single procentriole that eventually develops into the new daughter (Gönczy and Hatzopoulos, 2019). Precise control of the duplication process ensures that dividing cells enter mitosis with only two centrosomes, which then guide assembly of a bipolar spindle to facilitate accurate segregation of the duplicated genome.

Centrosome duplication is a complex, multi-step process that is triggered by the Ser/Thr kinase Polo-like Kinase 4 (Plk4) (ZYG-1 in *C. elegans*) (Zitouni et al., 2014). Plk4 is considered the master-regulator of centrosome duplication because its catalytic activity is required for centriole assembly and overexpression of Plk4 is sufficient to induce centriole overduplication (known as ‘centriole amplification’) (Bettencourt-Dias et al., 2005; Habedanck et al., 2005; Kleylein-Sohn et al., 2007). Moreover, Plk4 overexpression can promote *de novo* centriole assembly in acentriolar cells (Peel et al., 2007; Rodrigues-Martins et al., 2007). Due to its importance in centriole assembly, cells have evolved several regulatory mechanisms to restrict daughter centriole assembly to a single event per cell cycle and suppress ectopic Plk4 activity in order to prevent rampant centriole (and thus, centrosome) amplification (Chan, 2011; Ignacio et al., 2019; Sabat-Pośpiech et al., 2019).

Plk4 is regulated in a cell cycle-dependent manner. Although Plk4 mRNA levels peak during mitosis, both in cultured fly and mouse cells (Fode et al., 1996; Rogers et al., 2009), Plk4 is primarily regulated post-translationally. Plk4 extensively autophosphorylates a degron adjacent to its kinase domain, which is then recognized by the F-box protein Slimb, a substrate-binding module of the SCF E3 ubiquitin-ligase (Cunha-Ferreira et al., 2009; Rogers et al., 2009; Holland et al., 2010; Guderian et al., 2010; Brownlee et al., 2011; Cunha-Ferreira et al., 2013; Klebba et al., 2013). Thus, Plk4 activation promotes its own ubiquitin-mediated proteolysis to suppress centrosome amplification. Although endogenous Plk4 is near undetectable in most interphase cells – a consequence of its robust ability to autodestruct – Protein Phosphatase 2A (PP2A) stabilizes Plk4 during mitosis in *Drosophila* cells by counteracting Plk4 autophosphorylation and degradation (Brownlee et al., 2011). Plk4 localizes as a ring around each centriole wall during early mitosis, but then breaks symmetry during mitotic exit when it is remodeled into a single spot, an aggregate termed the ‘pre-procentriole’, from which the daughter centriole will later emerge (Dzhindzhev et al., 2017). Ring-to-spot remodeling of Plk4’s centriolar pattern also occurs in proliferating human cells, except this happens as cells approach the G1-to-S-phase transition, concurrent with procentriole assembly (Kim et al., 2013; Ohta et al., 2014; Park et al., 2014).

All Polo-like kinase family members contain an N-terminal kinase domain followed by one or more Polo box (PB) motifs (each ∼100 amino acids) separated by linkers of varying length (Zitouni et al., 2014) (Figure 1 – figure supplement 1A). Polo boxes function as hubs of protein interaction and adopt a characteristic Polo box fold, resembling half of a clamshell (Lowery et al., 2005). Plk4 is unique because it contains three distinct Polo boxes (PB1-3) while other Polo-like kinases contain only two (Slevin et al., 2012; Raab et al., 2021). Current models of Plk4 regulation suggest a stepwise pathway of activation that relies on its Polo boxes. Initially, a Plk4 molecule exists in an inactive, autoinhibited state mediated by its Linker 1 (L1), which connects the kinase domain to PB1 (Klebba et al., 2015a). L1 is thought to interact with the activation loop of the kinase domain and block its autophosphorylation, a critical step in Plk4 activation (Klebba et al., 2015a; Lopes et al., 2015). Relief of autoinhibition requires the C-terminal PB3 which forms a complex with the Plk4 activator, STIL (the human homolog of *Drosophila* Ana2), and likely repositions L1 through an interaction between Ana2/STIL and a short coiled-coil segment (CC) adjacent to the kinase domain (Arquint et al., 2015; Klebba et al., 2015a; Moyer et al., 2015; McLammarh et al., 2018). Subsequently, autophosphorylation of the Slimb-binding degron within L1 leads to polyubiquitation of several key lysine residues in PB1 (Klebba et al., 2015a).

Another unique and important feature of Plk4 is its ability to homodimerize. Studies of mouse Plk4 initially attributed homodimerization to PB3 but, interestingly, *Drosophila* PB3 is reportedly monomeric, whereas human PB3 can form monomers, dimers, and tetramers (Leung et al., 2002; Park et al., 2014; Arquint et al., 2015; Cottee et al., 2017). However, additional structural studies have revealed PB2, which resides within the central tandem Polo box cassette (PB1-PB2, also known as the ‘cryptic polo box’), as a conserved homodimerization module that adopts an end-to-end Z-shaped formation (Figure 1A) (Slevin et al., 2012; Park et al., 2014; Shimanovskaya et al., 2014). Since autophosphorylation of both the activation loop and Slimb-binding degron occurs in *trans* (Guderian et al., 2010; Klebba et al., 2013; Lopes et al., 2015), the purpose of homodimerization is believed to promote these important *trans*-autophosphorylation events. However, mutations in Plk4 that specifically prevent homodimerization do not exist and, thus, the function of homodimerization is yet unknown.

**Figure 1.**
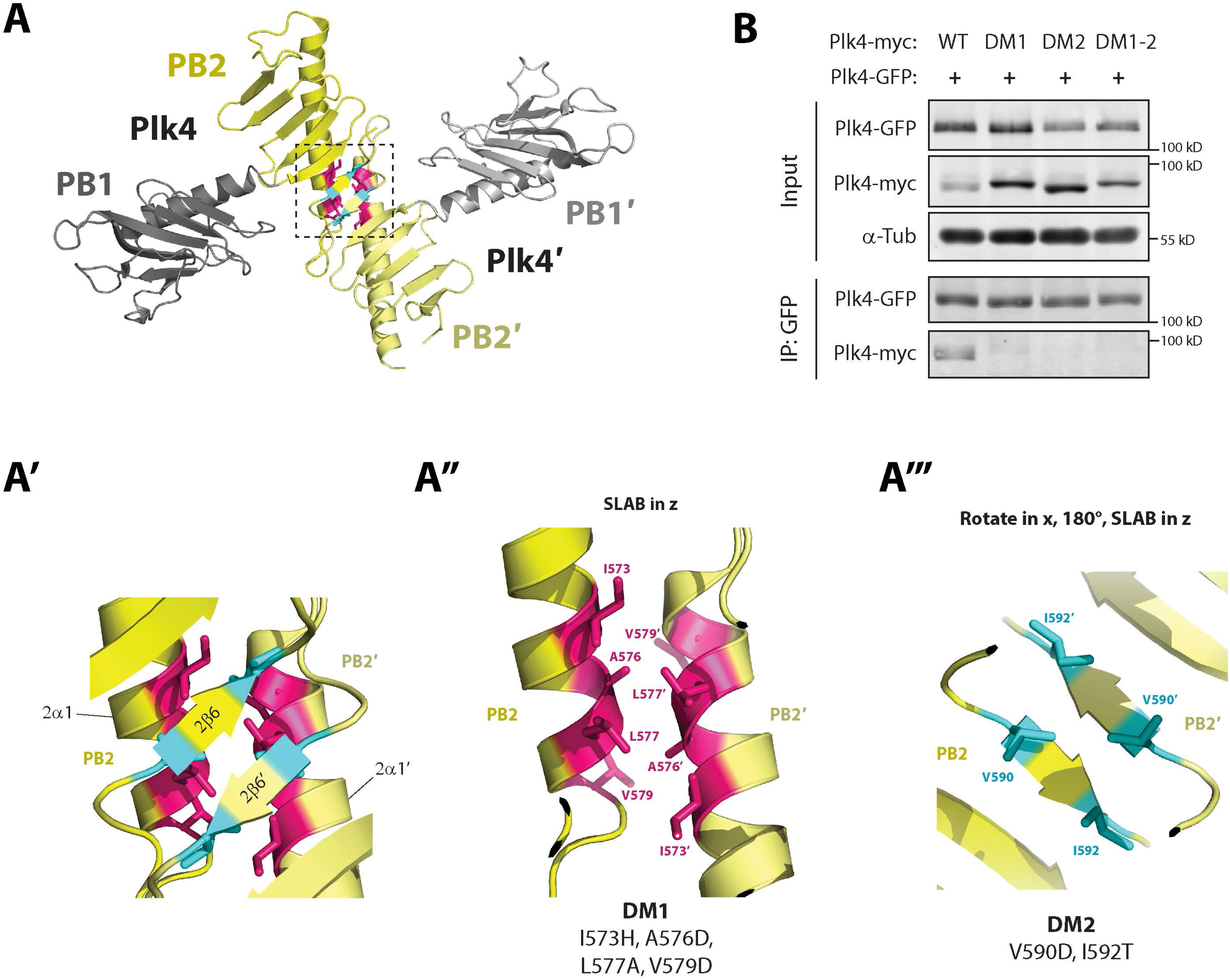
Generating a Plk4 dimerization mutant. (**A**) Ribbon diagram showing the quaternary structure of the *Drosophila* Plk4 PB1-PB2 which forms a Z-shaped end-to-end dimer. The dimerization interface (boxed) occurs between PB2-PB2’ (yellow). (**A’**) Magnification of boxed region in **A** highlighting the interaction interface. Six hydrophobic residues (red and blue) interact at the PB2-PB2’ interface. Residues responsible for dimerization are (**A’’**) four residues (red) on each 2α1 helix and (**A’’’**) two residues (blue) on each 2β6 strand. The specific amino acid substitutions that comprise the dimerization mutants DM1 and DM2 are listed. (**B**) Anti-GFP immunoprecipitates were prepared from lysates of S2 cells transiently co-overexpressing the indicated inducible Plk4-GFP and myc-tagged Plk4 constructs. Blots of the input lysates and IPs were probed for α-tubulin, GFP, and myc. Note that all three dimer mutants block Plk4 dimerization.

Notably, one face of the elongated PB1-PB2 Z-dimer contains a lengthy groove of basic residues that spans the tandem Polo boxes of each molecule (Park et al., 2014; Shimanovskaya et al., 2014). These basic patches act as docking sites for the acidic regions found in the N-terminus of SPD-2/Cep192 and Asterless (Asl)/Cep152, both of which are centriole-targeting factors for Plk4 (Cizmecioglu et al., 2010; Dzhindzhev et al., 2010; Hatch et al., 2010; Kim et al., 2013; Sonnen et al., 2013; Park et al., 2014; Shimanovskaya et al., 2014). In *Drosophila*, only Asl contains this acidic region, which makes several electrostatic contacts with patches of basic residues in PB1-PB2 (Park et al., 2014; Shimanovskaya et al., 2014; Klebba et al., 2015b).

Asl does more than simply act as a centriole receptor for Plk4, as depletion of Asl in cultured *Drosophila* S2 cells hinders Plk4’s ability to autodestruct, leading to increased Plk4 protein levels (Klebba et al., 2015b). Asl also activates Plk4 kinase activity both *in vitro* and in cells (Boese et al., 2018). However, the mechanisms by which Asl activates Plk4 remain a mystery. Since Asl binds PB1-PB2, one possibility is that Asl brings the dimerization interfaces of two Plk4 molecules together, facilitating both homodimerization and *trans*-activation. Here, we examine the role of homodimerization in Plk4 activity by generating and analyzing separation-of-function mutations in fly Plk4 that block its homodimerization. Surprisingly, our findings demonstrate that Plk4 homodimerization is not required for efficient activation or function of the kinase, and that Plk4 can be activated by an alternative activation pathway involving its aggregation.

## Results

### *Drosophila* Plk4 Polo Box 3 (PB3) is monomeric

To determine homodimerization’s in regulating Plk4 activity, we initially sought to identify regions of *D. melanogaster* (Dm) Plk4 required for oligomerization. We first determined the oligomeric state of purified DmPlk4 PB3 by solving a 1.75 Å crystal structure of this domain (Figure 1 – figure supplement 1B; table supplement 1). DmPlk4 PB3 is monomeric and displays a canonical Polo box fold: a six-stranded β-sheet with an α-helix laying across and orthogonal to the β-strands. Size exclusion chromatography with multiangle light scattering (SEC-MALS) analysis confirmed the monomeric state of DmPlk4 PB3 in solution (Figure 1 – figure supplement 1C). Our findings agree with prior work that found DmPlk4 PB3 to be a monomeric, canonical Polo box domain (Cottee et al., 2017). In contrast, SEC-MALS analysis of *Homo sapiens* (Hs) Plk4 PB3 revealed a dimeric solution state (Figure 1 – figure supplement 1C). While this contrasts with some reports describing HsPlk4 PB3 as a monomer (Park et al., 2014; Arquint et al., 2015), these results support studies describing mammalian (mouse and human) Plk4 PB3 to be oligomeric; a strand-swapped dimer capable of forming transient tetramers (i.e., dimers of strand-swapped dimers) (Leung et al., 2002; Cottee et al., 2017). Thus, our data suggest that PB3 does not contribute to fly Plk4 homodimerization, consistent with our previous co-immunoprecipitation experiments showing that Plk4 constructs lacking PB3 can still dimerize (Klebba et al., 2015a). Moreover, HsPlk4 PB3 may transition between monomeric and oligomeric states but the functional relevance of this is unknown.

### Mutation of residues within the PB2-PB2 interface disrupt Plk4 dimerization

Crystal structures of the tandem Polo box cassette (PB1-PB2) have revealed a variety of Plk4 dimer configurations with the Z-shaped homodimer being the most common, having been observed in *C. elegans*, *Drosophila*, and human Plk4 (Slevin et al. 2012; Park et al., 2014; Shimanovskaya et al., 2014) (Figure 1A, A’). Within *Drosophila* Plk4, Z-dimers form with a symmetric PB2-PB2 interface that is stabilized by hydrophobic interactions involving four residues in 2α1 helix (Figure 1A’, red) and two residues in 2β6 strand (Figure 1A’, blue). Based on this structure, we made two sets of amino acid substitutions designed to specifically ablate homodimerization: Dimer Mutant 1 (DM1) that altered all four 2α1 helix residues (I573H/A576D/L577A/V579D) (Figure 1A’’) and DM2 that changed two residues in 2β6 (V590D/I592T) (Figure 1A’’’), as well as a mutant (DM1-2) containing all 6 substitutions. Substitutions were chosen that either introduce electrostatic repulsive charges or diminish contacts at the PB2-PB2 interface, while maintaining stabilizing interactions within the Polo box core.

To test whether these mutations prevent oligomerization, we performed dimerization assays using cultured *Drosophila* S2 cells co-expressing wild-type (WT) Plk4-GFP with different myc-tagged Plk4 constructs. WT Plk4-GFP protein was immunoprecipitated from clarified cell lysates and immunoblotted for associated Plk4-myc. As expected, WT Plk4-myc bound WT Plk4-GFP (Figure 1B**, lane 1**). Strikingly, introducing any of the dimerization mutations in Plk4-myc blocked its ability to co-immunoprecipitate with WT Plk4-GFP (Figure 1B**, lanes 2-4**). We next examined how dimerization mutations in PB2 influence its proper folding *in silico* using the program AlphaFold which predicts protein structures (Jumper et al. 2021). Whereas PB2 harboring DM1 or DM2 adopted a proper Polo box fold, DM1-2 was predicted to miss the 2α6 strand (Figure 1 – figure supplement 2). Therefore, DM2 was chosen for most of our study because it contained the least number of substitutions and superimposed seamlessly with WT PB2 (Figure 1 – figure supplement 2B).

### Plk4 homodimerization is required to generate an Asl-binding site

To examine the impact of homodimerization on Plk4 activation, we first utilized an *in vitro* approach by purifying full-length WT or DM2 Plk4 proteins, mixing them with γ^32^-ATP, and measuring their ability to autophosphorylate, as Plk4 itself is a physiological substrate (Figure 2A; first treatment in graph). We found that WT Plk4 activity was low, presumably because the majority of the kinase exists in an autoinhibited state, as previously described (Boese et al., 2018). Although DM2 possessed lower activity than WT, this difference was not significant. Our results suggest that the simple act of dimerization is not sufficient to relieve autoinhibition.

**Figure 2.**
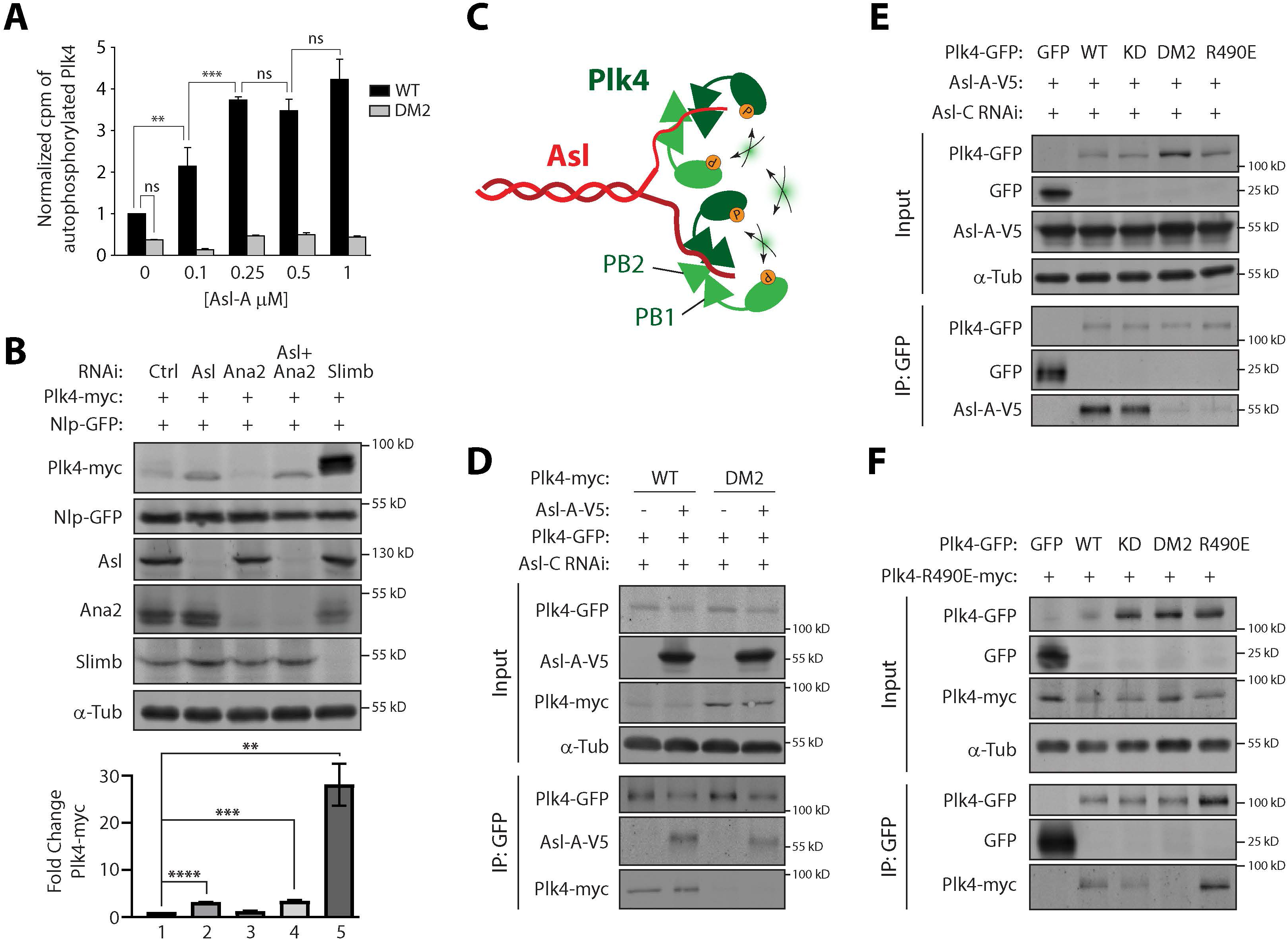
Plk4 homodimerization does not stimulate kinase activation but, instead, functions to generate an Asl-A binding site. (**A**) Asl-A stimulates WT Plk4 kinase activity but not the DM2 dimer mutant. Purified full-length WT or DM2 Plk4 (0.25 μM) was incubated with the indicated concentrations of purified Asl-A and γ^32^-ATP. The Plk4 in each sample (resolved by SDS-PAGE) was scintillation counted (cpm) and then normalized to the count of the WT/no AslA sample in that experiment. There was no significant differences in activity between any of the DM2 samples, regardless of Asl-A concentration. Asterisks mark significant differences as determined by one-way ANOVA test with Tukey’s multiple comparisons post-test. *n* = 3 experiments. Error bars, SEM. For all figures, *, 0.05 ˃ P ≥ 0.01; **, 0.01 ˃ P ≥ 0.001; ***, 0.001 ˃ P ≥ 0.0001; ****, 0.0001 ˃ P; ns, not significant. (**B**) Depletion of Plk4 activator, Asl, but not Ana2, increases Plk4 levels. (*Upper panels*) S2 cells were RNAi treated for 7 days. On day 4, cells were co-transfected with Nlp-GFP (transfection loading control) and inducible Plk4-myc. Starting day 5, cells were induced to express Plk4-myc. Cell lysates were prepared on day 7 and probed on immunoblots for GFP, myc, Asl, Ana2, and α-tubulin (loading control). (*Graph*) Shows relative amounts of Plk4-myc as determined by densitometry, normalized to Nlp-GFP, and plotted relative to control RNAi. *N* = 3 experiments. Asterisks mark significant differences from two-tailed t-tests comparing each condition to control. *n* = 3 experiment per treatment. Error bars, SEM. (**C**) Schematic showing how dimeric Asl-A (red) could facilitate *trans*-autophosphorylation of Plk4 (green) by promoting homodimerization of the kinase as well as positioning two dimers in close proximity. For simplicity, PB3 of Plk4 is not shown. (**D**) Asl-A does not promote the dimerization of a Plk4 dimer mutant. S2 cells were Asl-depleted by RNAi targeting an exon in the gene region encoding Asl-C for 7 days. On day 4, cells were transfected with the indicated constructs and starting day 5, induced to express for 48 hours. Anti-GFP immunoprecipitates (IPs) were prepared from lysates and Western blots of the IPs were probed for α-tubulin, GFP, V5, and myc. Note that DM2 Plk4-myc fails to immunoprecipitate with WT Plk4-GFP in cells co-expressing Asl-A-V5 (lane 4). (**E**) Plk4 dimerization is required for Asl-A binding. Cells were prepared as described in **D**. Anti-GFP immunoprecipitates were prepared from lysates of Asl-depleted S2 cells transiently co-overexpressing the indicated inducible Plk4-EGFP and Asl-A-V5 constructs. Immunoblots of the input lysates and immunoprecipitates were probed for α-tubulin, GFP, and V5. Note that Asl-A binding is negligible to DM2 (lane 4) as well as the control Asl-binding mutant, R490E Plk4 (lane 5). (**F**) The Asl-binding mutation R490E does not prevent Plk4 dimerization. Anti-GFP immunoprecipitates were prepared from lysates of S2 cells transiently co-overexpressing the indicated inducible Plk4-EGFP and myc-tagged constructs. Blots of the input lysates and IPs were probed for α-tubulin, GFP, and myc.

We next examined how Plk4 homodimerization might affect an interaction with its regulators. Notably, Plk4 kinase activity is stimulated by ‘activators’ including human STIL and *Drosophila* Asl (Arquint et al., 2015; Moyer et al., 2015; Boese et al., 2018). Depletion of STIL in cultured human cells or Asl in S2 cells increases Plk4 protein levels because Plk4’s ability to autophosphorylate, and hence autodestruct, is impaired (Klebba et al., 2015b; Moyer et al., 2015). Therefore, we depleted either Asl, Ana2 (the fly homolog of STIL), or both using RNAi in S2 cells and then measured the levels of exogenous Plk4. Whereas depletion of Ana2 had no effect on Plk4 levels (Figure 2B, lane 3), Asl RNAi elevated Plk4 levels and this did not further increase in double-depleted cells (Figure 2B, lanes 2 and 4). Depletion of the F-box protein Slimb, which is responsible for Plk4’s ubiquitination (Cunha-Ferreira et al., 2009; Rogers et al., 2009), caused a dramatic accumulation of Plk4 (Figure 2B, lane 5). These findings demonstrate that Asl is an important Plk4 activator in fly cells, unlike human cells that rely on STIL to suppress global Plk4 levels (Moyer et al., 2015). Intriguingly, Plk4 levels are much higher in Slimb-depleted cells than Asl-depleted cells, indicating the presence of an Asl-independent mechanism that eliminates the vast majority of transgenic Plk4 in these cells.

Asl contains large regions of predicted coiled-coil and can be divided into three scaffolding units, which we refer to as Asl-A, -B and -C (Figure 2 – figure supplement 1A) (Varmark et al., 2007; Dzhindzhev et al., 2010; Klebba et al., 2015b). Asl-A is a Plk4-binding module known to regulate Plk4 activity (Klebba et al., 2015b, Boese et al., 2018). The N-terminus of Asl-A contains the conserved acidic region (30 amino acids) that binds along one face of the PB1-PB2 Z-dimer (Park et al., 2014; Shimanovskaya et al., 2014) (Figure 2 – figure supplement 1B). We also mapped this interaction by charge switching the two most conserved acidic residues in this region to lysine (E24K/E25K), which we found abolished Plk4 binding by co-immunoprecipitation (Figure 2 – figure supplement 1C). Asl-B acts as an extended linker between the functional Asl-A and Asl-C domains. Asl-C functions as a platform capable of binding several proteins that localize to the centriole surface (Galletta et al., 2016). Importantly, Asl-C also contains a second Plk4-binding domain which, by itself, is sufficient to stabilize Plk4, target the kinase to centrioles, and induce centriole assembly (Klebba et al., 2015b). Whereas Asl-C is thought to mediate these functions during mitosis, Asl-A is functionally distinct from Asl-C; Asl-A globally activates Plk4 during interphase to promote Plk4 degradation, likely to prevent unwanted centriole amplification (Klebba et al., 2015a; Boese et al., 2018).

Previously, we reported that purified Asl-A robustly stimulates Plk4 catalytic activity *in vitro* (Boese et al., 2018). How Asl-A activates Plk4 is unclear, though. Notably, Asl-A can dimerize (Klebba et al., 2015b), and contains three segments of predicted coiled-coil (CC1-3) that could mediate its dimerization (Figure 2 – figure supplement 1A). Using Asl-A truncation and deletion mutants coupled with co-immunoprecipitation, we mapped the dimerization domain to CC3 (located within the C-terminal half of the protein opposite the acidic region) and confirmed that Asl-A monomers can bind Plk4 (Figure 2 – figure supplement 1D, E). Since Plk4 binds the acidic region with a stoichiometry of 2:1 (Plk4:Asl) (Park et al., 2014; Shimanovskaya et al., 2014), a single acidic region may enhance Plk4 homodimerization by bringing together two molecules of the kinase and promoting their *trans*-autophosphorylation (as proposed in Klebba et al., 2015b). Perhaps even greater activation may be achieved if dimeric Asl-A binds two Plk4 homodimers, placing them in close-proximity (see schematic in Figure 2C). To test this, we mixed full-length WT or DM2 Plk4 with an increasing amount of purified Asl-A and again measured Plk4 autophosphorylation *in vitro* (Figure 2A). As expected, addition of Asl-A stimulated WT autophosphorylation, reaching saturation at approximately a 2:1 (Plk4:Asl-A) stoichiometry. However, Asl-A failed to stimulate DM2 kinase activity. These results suggest that (1) Plk4’s innate ability to homodimerize is required in order for Asl-A to stimulate kinase activity and (2) Asl-A does not stimulate Plk4 activity by simply facilitating Plk4 oligomerization. To further test this second idea, we expressed Plk4-WT-GFP along with either WT or DM2-myc and asked whether co-expression of Asl-A could promote Plk4 dimerization. Because Asl oligomerizes, we eliminated the influence of endogenous Asl on the binding of the Asl-A fragment by first depleting S2 cells of Asl using RNAi targeting the Asl-C coding region. As expected, Plk4-WT-GFP co-immunoprecipitated both Plk4-WT-myc and Asl-A when co-expressed (Figure 2D, lanes 1 and 2). However, DM2-myc did not co-immunoprecipitate with Plk4-WT-GFP (Figure 2D, lanes 3) even when it was co-expressed with Asl-A (Figure 2D, lane 4). These findings suggest that Asl-A is not sufficient to overcome the lack of affinity of the DM2 mutant for WT, and that Plk4 dimerization is likely mediated solely by the PB2-PB2 interaction.

One interpretation of our *in vitro* findings is that Plk4 homodimerization is a prerequisite for Asl-A binding. To test this, we co-expressed full-length Plk4-GFP (WT or mutant) and Asl-A in S2 cells depleted of endogenous Asl, then immunoprecipitated Plk4-GFP, and probed for Asl-A in the immunoprecipitate (Figure 2E). Asl-A associated with Plk4-WT as well as a kinase-dead (KD) mutant, demonstrating that this interaction is not dependent on kinase activity, consistent with previous findings (Klebba et al., 2013). Strikingly however, Asl-A failed to bind the DM2 mutant (Figure 2E, lane 4). In this assay, we also used a Plk4 mutant harboring the R490E substitution in PB1 to block binding to Asl-A, because the R490E substitution was previously shown to prevent binding between purified PB1-PB2 and a short Asl fragment encoding the acidic region (Shimanovskaya et al., 2014). As with DM2, Asl-A also failed to co-immunoprecipitate with Plk4 R490E (Figure 2E, lane 5). However, unlike DM2, the R490E Asl-A binding-mutant did not affect Plk4 dimerization: Plk4-R490E-myc co-immunoprecipitated with GFP-tagged WT, KD, and R490E Plk4, but not DM2 (Figure 2F), demonstrating that the substitutions within PB1 and PB2 produce clear separation-of-function phenotypes. Moreover, an AlphaFold-predicted structure of PB1-PB2 containing the R490E substitution is not misfolded, appearing indistinguishable from the WT protein (Figure 2 – figure supplement 2). Taken together, our data suggest a multistep pathway for Plk4 catalytic activation. Initially, monomeric Plk4 is autoinhibited and then homodimerizes through PB2. However, dimerization is not sufficient to relieve autoinhibition but functions to create a binding site for Asl, which then stimulates kinase activity by relieving autoinhibition.

### Plk4 homodimerization is not required for kinase activation or function

Thus far, our results predict that homodimerization should be an important step in the catalytic activation of Plk4 in cells. Accordingly, expression of a dimer mutant would be mostly inactive, and so its protein level would increase due to a failure in autodestruction. To test this prediction, we overexpressed Plk4 dimer mutants in S2 cells and compared their protein levels to WT and kinase-dead (KD) proteins (Figure 3A). (Endogenous Plk4 should have no influence on these experiments because its levels are nearly undetectable [Rogers et al., 2009] and dimer mutants are incapable of binding WT Plk4 [Figure 1B].) Plk4 levels were normalized to the KD level, which is maximal because of the KD mutant’s inability to autodestruct. As expected, WT levels were extremely low compared to KD (Figure 3A, lanes 1 and 2). Surprisingly however, levels of all Plk4 dimer mutants were intermediate (∼5x greater than WT and ∼4x less than KD levels) (Figure 3A, lanes 3-5), suggesting that Plk4 dimer mutants are mostly active to promote their degradation in cells. Near identical results were observed with the R490E mutant whose level also decreased significantly compared to KD (Figure 3B), indicating that Asl-A binding and stimulation are not strictly required for activation. In an alternative approach, we used RNAi to first deplete Asl and then measured the levels of different Plk4 proteins (Figure 3C). DM2 levels were unaffected by Asl depletion and dramatically reduced compared to KD (Figure 3C, lanes 5 and 6). Therefore, Plk4 can degraded by a process(es) that is independent of both Asl binding and homodimerization.

**Figure 3.**
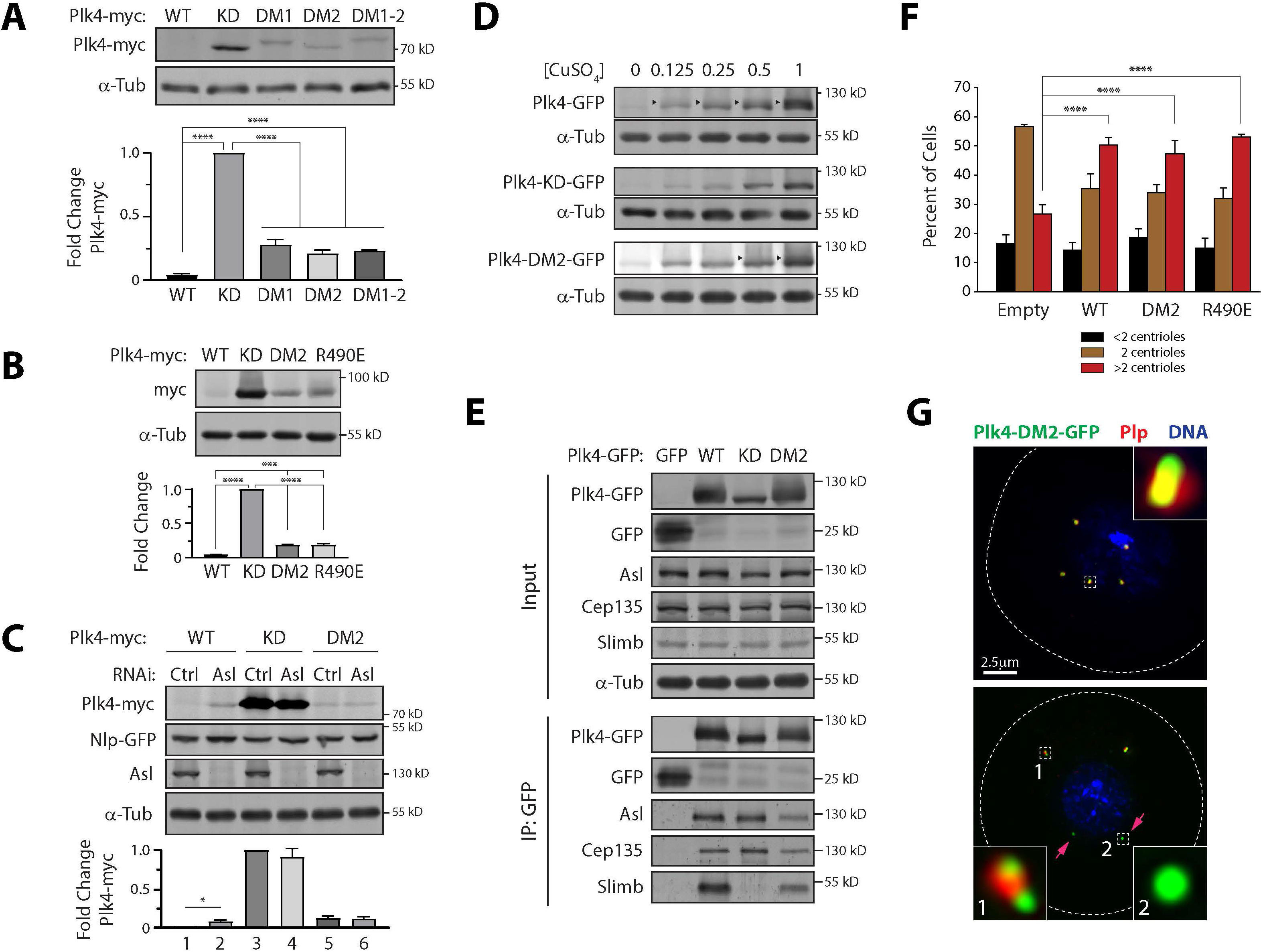
Dimer mutant Plk4 is catalytically active in cells and induces centriole amplification. (**A**) (*Upper panels*) Plk4 dimerization mutants efficiently autodestruct. The stabilities of different Plk4 proteins were analyzed by immunoblotting lysates of S2 cells transiently expressing the indicated inducible Plk4-myc construct. Anti-myc and α-tubulin (loading control) immunoblots are shown. (*Graph*) Plk4-myc intensities were measured by densitometry of the anti-myc immunoblots and normalized to their respective loading controls. Plotted values are the normalized Plk4 intensities relative to the kinase-dead (KD) sample. For all graphs in this figure, asterisks mark significant differences determined by one-way ANOVA with Tukey’s multiple comparisons post-test. *n* = 3 experiments. Error bars, SEM. (**B** and **C**) Asl is not required for Plk4 autodestruction. (*Upper panels*) The stabilities of different Plk4 proteins were analyzed by immunoblotting lysates of S2 cells transiently expressing the indicated inducible Plk4-myc constructs. Anti-myc, GFP, Asl, and α-tubulin (loading control) immunoblots are shown. (*Graphs*) Plk4-myc intensities were measured by densitometry of the anti-myc immunoblots and normalized to their respective loading controls. Plotted values are fold changes of the normalized Plk4 intensities relative to the kinase-dead (KD) sample. Nlp-GFP, loading control for transfected cells in **C**. For Asl RNAi, cells were treated as described in **2D**. *n* = 3 experiments. Error bars, SEM. (**D**) Dimerization mutant Plk4 can autophosphorylate. The electrophoretic mobilities of different Plk4 proteins were examined by immunoblotting lysates of S2 cells transiently expressing the indicated inducible Plk4-GFP constructs, using increasing concentrations (mM) of CuSO_4_ to drive increasing expression levels. Anti-GFP and α-tubulin (loading control) immunoblots are shown. Arrowheads mark slower-migrating species of phospho-Plk4. (**E**) Dimerization mutant Plk4 binds Slimb. S2 cells were transfected with the indicated inducible Plk4-GFP or GFP constructs and their expression induced with 1 mM CuSO_4_ for 24 hours. Anti-GFP IPs were prepared from cell lysates, and Western blots of the IPs were probed for GFP, Slimb, Asl, Cep135, and α-tubulin. (**F**) Plk4 homodimerization or Asl-A binding is not required to induce centriole amplification (>2 centrioles per cell). Transfected S2 cells were induced to express different Plk4-GFP constructs at low (0.25 mM CuSO_4_) levels for 3 days, then cells were fixed and immunostained for Plp (centriole marker) and their centrioles were manually counted. Centriole amplification occurs in cells expressing either wild-type (WT), DM2 or R490E Plk4 compared to cells transfected with control empty vector. *n* = 3 experiments. Error bars, SEM. (**G**) Plk4 homodimerization is not required for its localization to centrioles. S2 cells expressing Plk4-DM2-GFP (green) were immunostained for Plp (red) to mark centrioles. DNA, blue. Dashed lines show cell borders. Insets show higher magnifications of the boxed regions. (*Upper panel*) example of DM2-expressing cell containing centriole amplification. (*Lower panel*) Overexpressed DM2 not only localizes to centrioles but can also aggregate into round punctae (red arrows) that do not co-localize with the Plp centriole marker.

We also asked whether Ana2/STIL (the only other known Plk4 activator) may play a role in DM2 destruction. Plk4-DM2 levels were not affected by Ana2 depletion (Figure 3 – figure supplement 1). Taken together, our findings are unexpected and suggest that Plk4 dimerization is not a prerequisite for relief of autoinhibition and efficient degradation. Equally surprising is our finding that the bulk of Plk4 degradation is independent of Asl binding-induced activation. Based on the level of WT Plk4 in Asl-depleted cells or DM2 compared to KD, Asl is responsible for stimulating the autodestruction of ∼12-20% of the total transgenic Plk4 in these cells. These findings reinforce our previous Slimb RNAi data (Figure 2B), suggesting that cells contain an unidentified pathway of Plk4 activation.

Since Plk4 extensively autophosphorylates to generate a phosphodegron for Slimb binding (amino acids 292-297) (Figure 1 – figure supplement 1A), a prerequisite step in its ubiquitination, we next examined the dimer mutant’s ability to autophosphorylate by assessing its mobility on SDS-PAGE. Induction of increasing levels of expressed Plk4-WT-GFP in cells clearly revealed the autophosphorylated, slower-migrating species of the kinase which was not observed with Plk4-KD, as previously shown (Klebba et al., 2015a; Klebba et al., 2015b) (Figure 3D, upper and middle panels). A similar phospho-species was observed for DM2 when the mutant was highly expressed (Figure 3D, lower panel). Moreover, we examined DM2’s ability to bind Slimb, an established proxy for kinase activity (Cunha-Ferreira et al., 2013; Klebba et al., 2013). As expected, Slimb failed to co-immunoprecipitate with Plk4-KD. In contrast, Slimb bound both Plk4-WT and DM2 (Figure 3E). These results further support the conclusion that dimer mutant Plk4 is catalytically active, promoting its autophosphorylation and Slimb-mediated degradation.

We next examined the ability of dimer mutant Plk4 to induce centriole assembly. Overexpression of Plk4-WT is sufficient to drive centriole amplification in cells and, at relatively low expression for 3 days (similar to Figure 3A), we measured a significant increase in the percentage of cells (∼2x) with an abnormally high number of centrioles (>2) compared to control cells transfected with empty vector (Figure 3F, red bars). Strikingly, expression of either DM2 or R490E induced a similar and significant increase in the frequency of centriole amplification (Figure 3F). These results suggest that neither homodimerization nor Asl binding are necessary for over-expressed Plk4 to induce centriole amplification.

Since DM Plk4 can promote centriole amplification, we expect that it should also localize to centrioles. Indeed, we found that Plk4-DM2-GFP targets centrioles as revealed by its co-localization with Plp, a protein that decorates the outer-surface of mature centrioles (Figure 3G) (Martinez-Campos et al., 2004). Importantly, Asl is a Plk4 centriole-targeting factor, and we previously discovered this targeting is mediated by an interaction between the PB1-PB2 cassette and a Plk4-binding domain within the C-terminus of Asl (called Asl-C) (Klebba et al., 2015b). Although DM2 cannot bind Asl-A (which relies on Plk4 dimerization [Figure 2E]), we found that DM2 binds endogenous Asl (Figure 3E), presumably through its association with Asl-C. These findings explain how dimer mutant Plk4 can still localize to centrioles and suggests that, unlike the acidic region in Asl-A, Plk4 dimerization is not required for Asl-C binding or centriole targeting. Moreover, Plk4 dimerization is not required to bind the centriole protein, Cep135 (Figure 3E), an interaction that also occurs via the PB1-PB2 cassette in Plk4 (Galletta et all., 2016).

### Dimerization is not required for Plk4 *trans*-activation or *trans*-phosphorylation

Phosphorylation of both the activation loop and the Slimb-binding phosphodegron within Plk4 occur via *trans*-autophosphorylation (Guderian et al., 2010; Klebba et al., 2013; Lopes et al., 2015). These important post-translational modifications have been thought to require Plk4 dimerization, and this notion is supported by experiments where two different Plk4 constructs are co-expressed in cells (Figure 4A,B), allowing them to interact, *trans*-phosphorylate and, thus, potentially influence the degradation of the other. The relative levels of transgenic Plk4 in cell lysates are then used as measures of relative kinase activities. For example, co-expression of Plk4-KD-myc with Plk4-WT-GFP leads to the *trans*-autophosphorylation of the degron in Plk4-KD and its degradation (Figure 4A, lane 2; **4B**, lane 6). Our current findings challenge the idea that homodimerization is the sole mechanism underlying *trans*-phosphorylation. To test this further, we co-expressed Plk4-DM2 with Plk4-KD (differently tagged) and found that, as with WT, DM2 promoted the destruction of KD (Figure 4A, lanes 2, 4; Figure 4B, lanes 5, 8). Similarly, co-expression of the Asl-binding mutant, Plk4-R490E, with Plk4-KD demonstrated that Asl binding is not required for *trans*-phosphorylation (Figure 4A, lane 5). Furthermore, co-expression of Plk4-DM2 with Plk4-WT did not reduce the level of DM2 compared to expression of DM2 alone (Figure 4B, lanes 7 versus lane 9), suggesting that, on its own, the intrinsic kinase activity of DM2 is sufficient to initiate Asl-independent degradation. Therefore, neither Plk4 homodimerization nor Asl-A binding is required for this *trans*-phosphorylation-induced degradation pathway.

**Figure 4.**
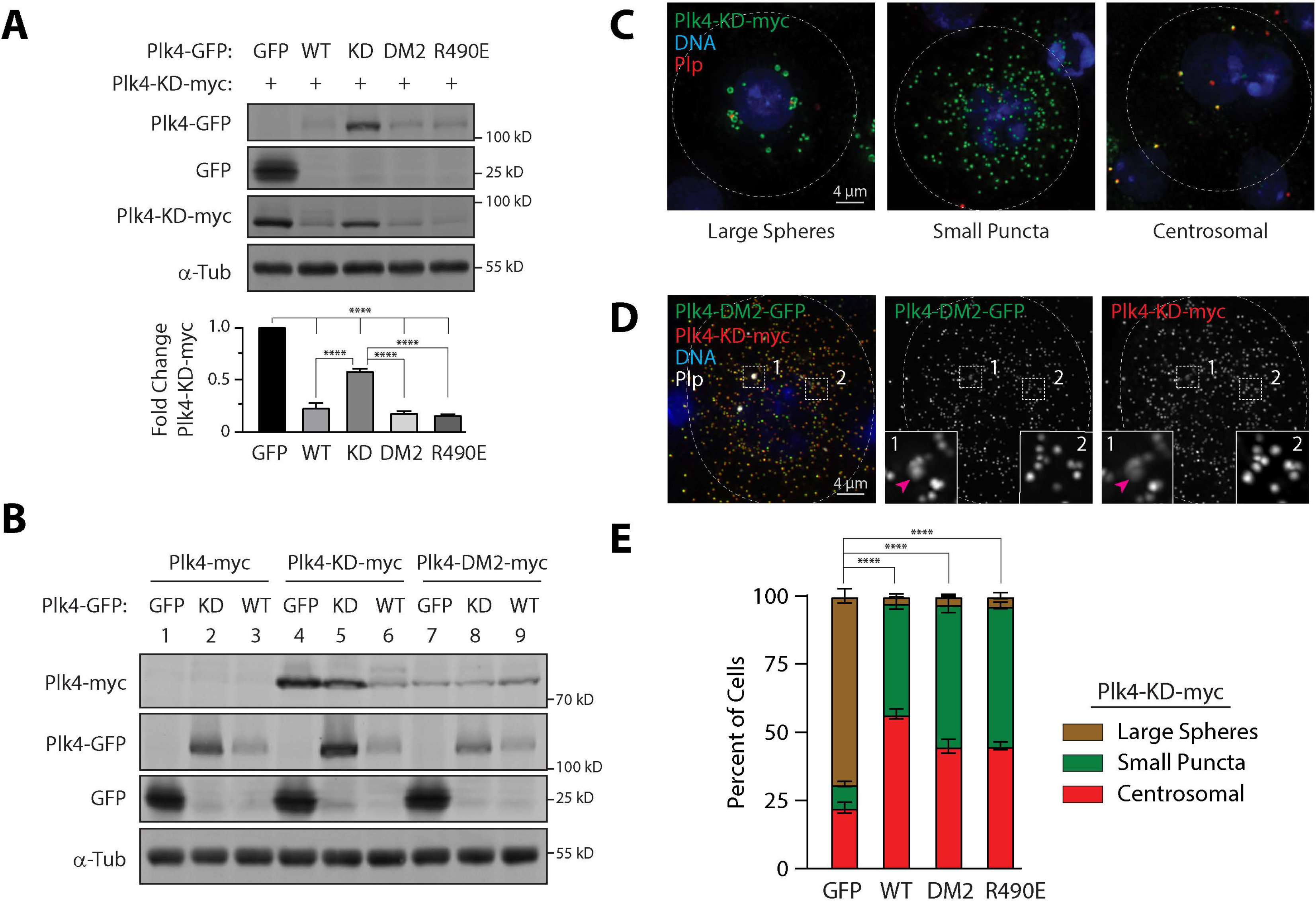
Plk4 can phosphorylate *in trans* to promote its degradation without having to homodimerize. (**A**) Plk4 *trans*-autophosphorylation-mediated degradation does not require Asl-A to stimulate kinase activity. (*Upper panels*) The stabilities of different Plk4 proteins were analyzed by immunoblotting lysates of S2 cells transiently expressing the indicated inducible GFP (control) or Plk4-GFP constructs along with Plk4-KD-myc. Anti-myc, GFP, and α-tubulin (loading control) immunoblots are shown. (*Graph*) Plk4-KD-myc intensities were measured by densitometry of the anti-myc immunoblots and normalized to their respective loading controls. Plotted values are the normalized Plk4-KD-myc intensities relative to the control GFP co-transfection treatment. Asterisks mark significant differences determined by one-way ANOVA with Tukey’s multiple comparisons test. Error bars, SEM. (**B**) WT Plk4 does not appreciably *trans* activate the Plk4 dimerization mutant. The stabilities of different Plk4 proteins were analyzed by immunoblotting lysates of S2 cells transiently expressing the indicated inducible GFP (control) or Plk4-GFP and -myc constructs. Anti-myc, GFP, and α-tubulin (loading control) immunoblots are shown. (**C-E**) Co-expression of dimer mutant or Asl-A-binding mutant Plk4 alters the pattern of Plk4 KD aggregates in S2 cells. (**C**) Images show S2 cells expressing Plk4-KD-myc (green) whose localization fits three distinct patterns. After 3 days of expression, cells were stained for Plp-labeled centrioles (red) and DNA (blue). Dashed yellow lines mark cell borders in all images. (**D**) DM2 and KD Plk4 co-aggregate. S2 cells expressed inducible Plk4 constructs for 3 days and were stained for Plk4-DM2-GFP (green), Plk4-KD-myc (red), centriole marker Plp (white), and DNA (blue). Dashed boxes are shown at higher magnification (insets). Arrowhead (pink) marks a centrosome. (**E**) Graph shows percentage of cells with the indicated Plk4-KD-GFP localization patterns. Note that all Plk4 constructs prevent KD Plk4 from forming large aggregates. Significant differences were determined using one-way ANOVA with Tukey’s multiple comparison test. *n* = 3 experiment per treatment. Error bars, SEM.

How could Plk4 *trans*-phosphorylation occur then if not through homodimerization? One possibility is that Plk4 aggregation might trigger its *trans*-activation. Indeed, a previous study found that local accumulation of Plk4 is sufficient to activate the kinase and promote *trans*-autophosphorylation of its activation loop (Lopes et al., 2015). This was demonstrated, in part, by targeting Plk4 to the lumen of peroxisomes where it was active without cytoplasmic Plk4 activators (such as STIL) (Lopes et al., 2015). Consistent with this hypothesis, we noticed small spherical aggregates of Plk4-DM2 that formed in the cytoplasm of some cells and that did not co-localize with centrioles (Figure 3G, bottom panel, pink arrows). These aggregates are highly reminiscent of Plk4 condensates observed by others (Montenegro Gouveia et al., 2018; Yamamoto and Kitagawa, 2019).

To test whether Plk4 could *trans*-phosphorylate through aggregation, we took advantage of Plk4-KD’s tendency to form numerous aggregates when expressed in cells (Klebba et al., 2013). Although it always localized to centrosomes, Plk4-KD displayed three distinct patterns when overexpressed in cells: 1) large spheres, 2) small punctae, and 3) mostly centrosomal (Figure 4C). Co-expression of Plk4-KD with other Plk4 constructs showed extensive co-localization of the two differently-tagged Plk4 proteins in these aggregates, including dimer mutant Plk4-DM2 (Figure 4D). We reasoned that if Plk4 was active in these aggregates to *trans*-phosphorylate KD protein and promote its degradation, then co-expression should reduce the frequency of cells that display Plk4-KD aggregates. After 3 days of co-expression, cells were stained and scored for their KD patterns. We found that most cells (∼70%) formed large spheres of Plk4-KD-myc when co-expressed with control GFP (Figure 4E), consistent with our Western blots of whole cell lysates showing high levels of the KD protein (Figure 4A,B). Strikingly however, co-expression of WT, DM2 or R490E mutant Plk4 significantly reduced the number of cells containing large spheres of KD, replacing them with the small punctae or mostly centrosomal patterns (Figure 4E). Taken together, our findings suggest that Plk4 *trans*-activation can occur via simple aggregation. Active forms of Plk4 can *trans*-phosphorylate KD within aggregates to promote its degradation, without homodimerization or Asl binding.

## Discussion

Plk4 is a unique member of the Polo-like kinase family because it contains three Polo boxes (PB1-3) and has the ability to homodimerize. Other Plks possess only two Polo boxes which can form intramolecular dimers but are otherwise monomeric (Zitouni et al., 2014). In this study, we sought to answer the question -- what is the purpose of Plk4 homodimerization? Our findings pinpoint a cluster of hydrophobic residues in *Drosophila* PB2 that mediate formation of the Z-shaped Plk4 homodimer, mutation of which selectively disrupts dimerization. With these separation-of-function mutations in hand, we determined that the function of dimerization is to create a binding site for the acidic region of Asl, located in the N-terminus of Asl (Asl-A), and that binds with a stoichiometry of 2:1 (Plk4:Asl-A) (Park et al., 2014; Shimanovskaya et al., 2014). Asl-A binding then relieves Plk4 autoinhibition and stimulates kinase activity. How binding of the acidic region in Asl-A relieves Plk4 autoinhibition remains an important unanswered question which will require a deeper molecular understanding of the mechanism of autoinhibition. Current models suggest that linker 1 (L1) interacts with the kinase domain to prevent Plk4 from *trans*-autophosphorylating three key residues in the activation loop (Klebba et al., 2015a; Lopes et al., 2015; Boese et al., 2018). Possibly, Asl-A binding to the nearby PB1-PB2 dimer repositions L1 and allows access.

The purpose of Plk4 homodimerization was thought to be prerequisite for *trans*-autophosphorylation of both the activation loop and the downstream regulatory element (DRE) (that becomes the Slimb-binding phosphodegron). Remarkably, our characterization of dimer mutant Plk4 suggests that dimerization is not required for *trans*-phosphorylating both activation loop and DRE. This is likely due to Plk4’s innate ability to aggregate, probably forming liquid condensates (Montenegro Gouveia et al., 2018; Yamamoto and Kitagawa, 2019). Indeed, Plk4 *trans*-activates in a concentration dependent manner (Lopes et al., 2015) and independently of activators, due to its inherently low catalytic activity (Klebba et al., 2015a). Conceivably, upon aggregation, Plk4 concentration reaches a critical mass where it is able to *trans*-phosphorylate the L1 domain, which then blocks further autoinhibition (Klebba et al., 2015a). Consequently, Plk4 kinase activity is dramatically stimulated owing to this positive feedback loop and, at high enough levels, can also induce the formation of supernumerary centrioles, as observed for WT or Asl-A binding mutant Plk4.

Although our work with dimer mutant Plk4 demonstrates that oligomerization is not required for its activation or function, we do not suggest that homodimerization is dispensable for proper Plk4 regulation. In fact, Plk4 homodimerization and the resulting *trans*-autophosphorylation that Asl-A would stimulate are important steps in a pathway of Plk4 autodestruction. Indeed, while mutants that cannot bind the acidic region in Asl and/or dimerize are efficiently degraded, their protein levels are higher compared to WT Plk4 (Figure 2B, 3A, 3C). In proliferating *Drosophila* cells, Plk4’s role in the centriole duplication cycle is confined to a small window of time during mitotic exit; during most of the cell cycle, Plk4 is not present due its robust ability to autodestruct (Rogers et al., 2009; Holland et al., 2010). Because endogenous levels of Plk4 are extremely low and near undetectable, homodimerization would immediately trigger Asl binding (a far more abundant protein) and *trans*-autophosphorylation would ensue. We propose that the Asl-mediated pathway of Plk4 activation maintains a low level of Plk4 and, thus, suppresses centriole amplification (Figure 5A-C).

**Figure 5.**
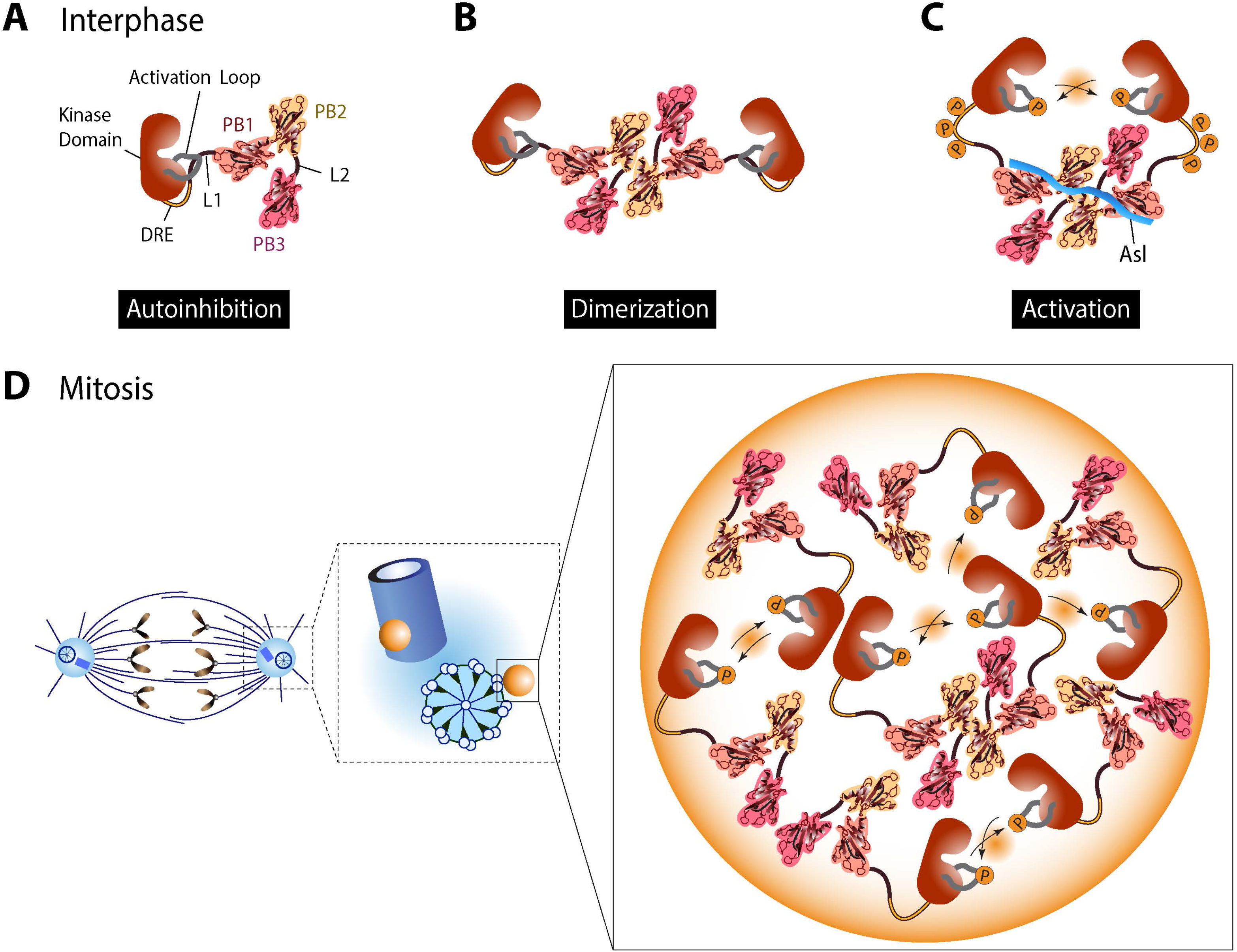
Multistep model describing cell cycle-dependent pathways of Plk4 activation. (**A**) In interphase cells, Plk4 is globally maintained at a low level due to an Asl-dependent pathway of Plk4 activation and autodestruction. Newly synthesized Plk4 initially exists in an autoinhibited, monomeric form. Autoinhibition is exerted by Linker 1 (L1) which binds to the activation loop of the kinase domain and prevents its autophosphorylation. (**B**) Plk4 homodimerizes through hydrophobic interactions between two PB2 modules. The resultant Z-shaped end-to-end dimer creates a unique Asl binding site that extends across the dimer and makes multiple electrostatic contacts across both PB1 and PB2. (**C**) The short acidic region within the N-terminus of Asl binds along one Plk4 Z-dimer surface and relieves autoinhibition by repositioning L1. Plk4 catalytic activity is dramatically activated by Asl binding (Boese et al., 2018; Figure 2). Plk4 then *trans*-autophosphorylates its activation loop as well as the Slimb-binding degron which promotes its ubiquitin-mediated proteolysis and, consequently, suppressing centriole duplication. (**D**) During mitosis, Plk4 is dephosphorylated by PP2A, causing Plk4 protein levels to rise (Brownlee et al., 2011). Plk4 initially decorates mitotic centrioles as a ring but then remodels into a single asymmetric spot (the pre-procentriole) as cells exit mitosis (Dzhindzhev et al., 2017). Asl targets Plk4 to centrioles (Dzhindzhev et al., 2010), but unlike Plk4, the Asl ring that forms on mitotic centrioles does not remodel into the pre-procentriole and, thus, is not present to activate Plk4 in the aggregate (Fu et al., 2016). However, Plk4 has an intrinsically low level of catalytic activity (Klebba et al., 2015a; Lopes et al., 2015), and within this aggregate (likely a mixture of Plk4 monomers and dimers), the density of Plk4 is high enough to promote *trans*-autophosphorylation. Plk4 activation ultimately promotes procentriole assembly on this single site. Normally this aggregation pathway is not observed in interphase cells because Plk4 levels are too low to form aggregates (due to the Asl-mediated pathway) but the existence of this pathway comes into view in interphase cells force to overexpress Plk4.

Due to the inability to detect endogenous Plk4, most studies of Plk4 regulation require overexpression of transgenes that produce Plk4 levels that are orders of magnitude higher than the endogenous level. In the course of using this same approach, our work serendipitously revealed an alternative pathway of activation involving aggregation and *trans*-activation of Plk4 that does not require its dimerization or Asl-A binding. Indeed, previous studies describing *trans*-autophosphorylation (that naturally required the Plk4-overexpression methodology) were likely observing *trans*-phosphorylation of Plk4 within aggregates and attributed this to *trans*-autophosphorylation of dimeric Plk4 as an underlying mechanism (Guderian et al., 2010; Klebba et al., 2013; Lopes et al., 2015). We propose that this new aspect of Plk4 regulation (i.e., the aggregation pathway for Plk4 activation) could be utilized in the centriole duplication cycle (Figure 5D). During mitosis in *Drosophila* cells, Plk4 levels rise due to PP2A which counteracts Plk4 autophosphorylation (Brownlee et al., 2011). Asl targets the kinase to the centriole surface (Dzhindzhev et al., 2010), and Plk4 then remodels to form a single concentrated spot devoid of Asl/Cep152 called the pre-procentriole (Park et al., 2014; Dzhindzhev et al., 2017). Within the pre-procentriole, Plk4 is at a high local concentration and, as a mixture of monomers and dimers, can *trans*-activate. Notably, in this pathway, Plk4 need not be restricted to the centriole surface to induce centriole assembly; Plk4 can also induce *de novo* centriole formation but only at high concentration (Lopes et al., 2015). Future studies are necessary to establish the protein composition of the pre-procentriole and to determine the precise nature and sequence of molecular events required for this earliest phase of centriole assembly.

## Materials and methods

### Cloning and dsRNA generation

Plk4 point mutants were generated by PCR-based site-directed mutagenesis using Phusion polymerase (Thermo Fisher) and then cloned into the pMT expression vector, which contains a Cu-inducible metallothionein promoter. Control dsRNA, and dsRNA targeting either Asl or Ana2 was transcribed from DNA templates consisting of the T7 RNA polymerase promoter sequence (5’-TAATACGACTCACTA) fused to gene-specific (or control) sequences. *In vitro* dsRNA synthesis was prepared as previously described (McLamarrah et al., 2018).

### Cell culture, transfection, and RNAi

*Drosophila* S2 cells were cultured in Sf-900II SFM media (Life Technologies) as previously described (McLamarrah et al., 2018). dsRNA and DNA constructs were transfected into S2 cells by nucleofection (Nucleofector II, Amaxa), as previously described (McLamarrah et al., 2018). Cells were transfected with up to 2 µg of DNA. For RNAi treatments, cells were transfected with 40µg dsRNA on days 0 and 4. DNA constructs were transfected concurrent with dsRNA on day 4. Gene expression was induced with 0.25 mM CuSO_4_ for 48 hours unless stated otherwise.

### Antibody Generation

Ana2 antibodies were raised in rabbits against synthesized peptide [CQLDLENIRNQPKLL] (Pocono Rabbit Farm and Laboratories). To generate affinity-purified antibodies, peptide was conjugated to Affi-gel resin (Bio-Rad) according to manufacturer’s protocol. Serum was then incubated with the immobilized peptide on resin, the resin extensively washed, and then the resin eluted of anti-peptide antibodies using a brief incubation with low pH buffer.

### Cell lysates and Immunoblotting

To generate whole cell lysates for Western blot analysis, cells were lysed in extraction buffer (50 mM Tris-HCl, pH 7.2, 150 mM NaCl, 0.5% Triton X-100, 1 mM DTT, and 0.1 mM PMSF), extract protein concentrations were determined by Bradford, followed by addition of Laemmli sample buffer. Lysates were boiled for 5-10 min and stored at −20°C. Equal amounts of total protein of each whole cell lysate were resolved by SDS-PAGE, transferred onto nitrocellulose (Amersham), blocked in blocking buffer (5% milk in PBS, 0.1% Tween-20), sequentially probed with primary and secondary antibodies, and then scanned on a LiCor Odyssey CLx imager with Image Studio software (LiCor Biosciences). Primary antibodies used for Western blotting included rat anti-Cep135 (1:1000; McLamarrah et al., 2018), guinea pig anti-Slimb (1:1000; Brownlee et al., 2011), rat anti-Asl (1:1000; Boese et al., 2018), rabbit anti-Ana2 (this study), mouse anti-V5 monoclonal (1:3000, Life Technologies), mouse anti-myc (1:3000, Cell Signaling), mouse anti-GFP monoclonal JL8 (1:3000, Clontech), and mouse anti-α-tubulin monoclonal DM1A (1:3000, Sigma-Aldrich). Primary antibodies were diluted as indicated in blocking buffer. Host-specific IRDye 800CW secondary antibodies (LiCor Biosciences) were prepared according to the manufacturer’s instructions and used at 1:3000 dilution in blocking buffer.

### Immunoprecipitation assays

GFP-binding protein (GBP) (Rothbauer et al., 2008) was fused to the Fc domain of human IgG (pIg-Tail) (R&D Systems), tagged with His_6_ in pET28a (EMD Biosciences), expressed in *E. coli* and purified on HisPur Cobalt resin (Fisher) according to manufacturer’s instructions (Buster et al., 2013). Purified GBP was bound to magnetic Protein A Dynabeads (ThermoFisher), and then covalently-linked by incubation with 20 mM dimethyl pimelimidate in PBS, pH 8.3, for 2 hours at room temperature followed by quenching with 0.2 M ethanolamine, pH 8.3, for 1 hour at room temperature. GBP-coupled Dynabeads were stored in PBS, 0.1% Tween-20 at 4°C. Prior to use, beads were equilibrated in IP buffer (50 mM Tris, pH 7.2, 125 mM NaCl, 1 mM DTT, 0.5% Triton X-100, 1x SigmaFast protease inhibitors [Sigma], 0.1 mM PMSF, and 1 μg/mL SBTI). Transfected cells were harvested and lysed in IP buffer, the lysate concentrations determined by Bradford assay, and the lysates diluted to 5 mg/mL. Lysates were then clarified by centrifugation at 10Kxg, 5min, 4°C. Input samples were generated from the clarified lysates by addition of Laemmli sample buffer. For IPs, GBP-conjugated beads were rocked with clarified lysates for 30 min, 4°C, washed four times by resuspending beads in 1 ml IP buffer, transferred to a new tube during the final wash, then boiled in an equal volume of 2x Laemmli sample buffer. Inputs and IPs were then analyzed by Western blot.

### Protein Purification

His_6_-Plk4-PB3 (aa 657-745) was expressed in BL21(DE3) *E. coli* under kanamycin selection and induced with 100 μM IPTG for 16 hours at 20°C. Cells were harvested, resuspended in lysis buffer (25 mM Tris pH 8.0, 300 mM NaCl, 10 mM Imidazole, 0.1 % β-ME) supplemented with 1 mM PMSF, and sonicated to lyse. Lysate was clarified at 23Kxg, 45 min, 4°C, and supernatant applied to a column of Ni^2+^-NTA resin (Qiagen). After washing, Plk4 PB3 was eluted using a 250 ml, 10-300 mM imidazole gradient in lysis buffer. Fractions containing Plk4 were pooled, supplemented with 1 mM CaCl_2_ and His_6_-tag was removed by digesting for 12 hrs, 4°C, with 1 μg/ml bovine α-thrombin. Digested Plk4 was filtered through 1 ml benzamidine sepharose (Cytiva) to remove α-thrombin, and then exchanged into 100 ml of 25 mM HEPES, pH 7.0, 0.1% β-mercaptoethanol (β-ME). loaded onto an SP-sepharose column (Cytiva), and eluted over a 250 ml, 0-1 M NaCl gradient in 25 mM HEPES, pH 7.0, 0.1% β-ME. Fractions containing Plk4 PB3 were pooled, exchanged into protein storage solution (25 mM HEPES, pH 7.0, 100 nM NaCl, 0.1% β-ME), concentrated to 15 mg/ml, and stored in liquid nitrogen. SeMet-substituted protein was expressed in B834 *E. coli* with SeMet media (Leahy et al., 1994) and induced and purified following the procedure for native Plk4 PB3. Selenomethionine (SeMet)-substituted Plk4 proved soluble and yielded crystals that diffracted; however, these crystals provided a very weak anomalous signal, preventing phase information. We hypothesized that the single methionine residue in PB3 (M696) may reside in a flexible loop, consistent with both secondary structure prediction algorithms and lack of phase information from SeMet-substituted protein-derived crystals. Therefore, we systematically mutated several hydrophobic residues at the end of predicted secondary structure features to methionine (L675M, V692M, and V723M) to provide structured sites of SeMet incorporation. Following solubility tests and crystallization trials, only V692M yielded SeMet-substituted protein crystals. HsPlk4 PB3 (aa 884-970) was purified using the same protein expression vector and system and purification protocols, and concentrated to either 4.6 or 8 mg/mL in 25 mM HEPES, pH 7.5, 300 mM NaCl, 0.1% β-ME, 0.2 g/L NaN_3_.

### Crystallization, data collection, and structure determination

Plk4 PB3 and SeMet-substituted V692M Plk4 PB3 were crystallized using a mother liquor (1 ml) containing 32% PEG 4000, 200 mM Li_2_SO_4_, and 200 mM Tri, pH 8.5. Crystals formed from drops containing 2 μl of 15 mg/ml protein stock and 2 μl mother liquor. Native crystals were transferred to a cryoprotectant condition containing mother liquor supplemented with 20% ethylene glycol and flash-frozen in liquid nitrogen. SeMet V692M Plk4 crystals were transferred to MiTeGen LV CryoOil (Mitegen) and flash-frozen in liquid nitrogen. Diffraction data were collected at the Advanced Photon Source SER-CAT beamline 22-ID (Native, 1.00000 Å; V692M Se peak SAD data, 0.97926 Å). Crystals belonged to the space group P2_1_ with two molecules in the asymmetric unit. Data were processed and scaled using the HKL2000 suite (Otwinowski and Minor, 1997). The Phenix program suite (Adams et al., 2010) was used to find selenium sites, phase, build, and refine the SeMet V692M Plk4 PB3 structure against a MLHL target with reiterative building in Coot (Emsley et al., 2010). Refinement was monitored using 10% of the data randomly excluded from the refinement and used to calculate an R_free_ value (Brünger, 1992). The final SeMet V692M Plk4 PB3 model was refined to 1.75 Å resolution, and the most complete protomer used as a molecular replacement search model for the high-resolution native data set and found the two molecules in the asymmetric unit. The native Plk4 PB3 model was reiteratively built and refined against a ML target, yielding an R value of 19.2 and a R_free_ value of 23.3. The model includes two Plk4 PB3 protomers: chain A (residues 660-745) and chain B (residues 660-693, 699-743) and 118 water molecules. DmPB3 coordinates have been deposited in the Protein Data Bank under accession code 7RL3.

### Size Exclusion Chromatography and Multi-Angle Light Scattering (SEC-MALS)

DmPlk4 PB3 was purified without cleaving the N-terminal His_6_ tag, concentrated to either 10 or 27 mg/mL, and exchanged into running buffer (25 mM HEPES, pH 7.5, 300 mM NaCl, 0.1% β-ME, and 0.2 g/L NaN_3_). A Superdex 200 10/300 GL gel filtration column (Cytiva) was equilibrated in the running buffer, and 100µL samples protein were injected onto the column. Eluate was passed in tandem through a Wyatt DAWN HELEOS II light scattering instrument and a Wyatt Optilab rEX refractometer (Wyatt, 1993). The light scattering and refractive index data were used to calculate the weight-averaged molar mass of each peak using the Wyatt Astra V software program (Wyatt Technology). Two DmPlk4 PB3 and HsPlk4 PB3 runs were performed for each purification scheme, with two purifications for each construct, for a total of eight experiments.

### *In silico* structure modeling

Structures of Plk4 PB1-2 (aa 382-602) or PB2 (aa 501-602) or the designated mutants were predicted by inputting the primary sequence into AlphaFold Colab Notebook (https://colab.research.google.com/github/deepmind/alphafold/blob/main/notebooks/AlphaFold.i pynb) (Jumper et al., 2021). Each prediction model was then visualized and overlaid in Chimera (Pettersen et al., 2004).

### Kinase Assays

Asl-A-13A-His_6_ and full-length fly Plk4 (WT and DM2 mutant, both with N-terminal His_6_ and MBP [maltose binding protein] tags) were expressed in BL21(DE3)pLysS *E. coli*. The 13 alanine substitutions in Asl-A-13A were not random but replaced native residues known to be phosphorylated by Plk4; phospho-null Asl-A-13A dramatically stimulates Plk4 kinase activity (Boese et al., 2018). Bacterial pellets were resuspended in a lysis buffer, homogenized (EmulsiFlex-C3 homogenizer [Avestin]), and then clarified by centrifugation (200Kxg, 30min, 4°C). For Plk4 constructs, lysis buffer was PBS (137 mM NaCl, 2.7 mM KCl, 10 mM Na_2_HPO_4_, 1.8 mM KH_2_PO_4_, 0.5 mM MgCl_2_), pH 7.3, 0.1% TX-100, 100 mM arginine, 10 mM imidazole, 15% (by volume) glycerol, 1 mM EGTA, 1 µg/mL soybean trypsin inhibitor (SBTI), 0.1 mM PMSF, and SigmaFast protease inhibitor cocktail (Sigma). For AslA-13A, a modified lysis buffer was used consisting of PBS, pH 7.3, 20 mM imidazole, 1 mM DTT, and 0.1 mM PMSF; for homogenization, this buffer was supplemented with 0.1% TX-100, 1 µg/mL SBTI, and 0.25x SigmaFast protein inhibitor cocktail.

Clarified supernatants were rocked with Complete His-Tag resin (Roche), 30 min, 4°C and then gently pelleted (100xg, 5 min, 4°C). Resins with Plk4 were washed twice with 30 bed volumes of lysis buffer (without SBTI or EGTA, and with 0.1x of protease inhibitor cocktail) by rocking resin with buffer for 5 min, 4°C and then pelleting (100xg, 5 min, 4°C), for each wash. Plk4 was eluted with a step gradient of imidazole (30-200 mM final) in lysis buffer (without SBTI, EGTA, or protease inhibitor cocktail). Resin with Asl-A-13A was washed sequentially with 30 resin bed volumes of: (1) modified lysis buffer with 0.1% TX-100, (2) modified lysis buffer + 300 mM additional NaCl, and (3, 4) two washes with modified lysis buffer. For each wash, resin was rocked with buffer for 10 min, 4°C, and then the supernatant removed after resin was pelleted by centrifugation (100xg, 5 min, 4°C). Asl-A-13A was eluted with a step gradient of imidazole (40-150 mM) in modified lysis buffer.

Protein-containing elution fractions were identified by SDS-PAGE, and fractions were pooled, concentrated with a centrifugal filter (Amicon Ultra-15, Millipore), and then exchanged into kinase buffer (40 mM HEPES, pH 7.3, 150 mM NaCl, 5 mM MgCl_2_, 1 mM DTT, 15% glycerol), re-concentrated, and held at 4°C. Protein concentration of full-length construct was calculated by first measuring total protein concentration by Bradford assay, then measuring the percent purity of full-length protein using densitometry of coomassie-stained SDS-PAGE gels, and finally taking the product of these measurements to calclulate the concentration (mg/mL).

*In vitro* kinase assays were performed in 30 µL volumes with purified Plk4 +/- Asl-A-13A, 100 µM ATP (spiked with γP^32^-ATP), 10% glycerol, and varying quantities of purified GST (known to not be phosphorylated by Plk4; Brownlee et al., 2011) to maintain a constant total protein concentration in all assays. Assays were performed at 22°C in kinase buffer (see above). With the exception of ATP, all components of each assay were combined at least 5 min before the reaction was initiated. Reactions were initiated by addition of ATP (in kinase buffer), incubated for 1hr, and terminated by the addition of Laemmli sample buffer, heating to 100°C for ∼2 min, and then transfer to −20°C. To measure phosphorylation, samples were resolved by SDS-PAGE, and the gels coomassie-stained, dried, and then autoradiographed to detect phosphorylated proteins. Subsequently, individual Plk4 and dMBP bands were cut from the gels and analyzed by scintillation counting.

### Immunofluorescence microscopy

S2 cells were spread on concanavalin A-coated glass bottom plates and fixed in ice cold methanol for 12 min. Cells were washed with PBS, 0.1% Triton X-100 and blocked with slide blocking buffer (5% normal goat serum in PBS, 0.1% Triton X-100) for 30 min, room temperature. Primary antibodies were diluted in blocking buffer (rabbit anti-Plp [1:3000], mouse anti-myc [1:3000]), and slides were incubated for 1 hour, room temperature. After 3 washes with PBS, 0.1% Triton X-100, cells were incubated at room temperature with Hoechst 33342 (Life Technologies, 3.2 μM final concentration) and secondary antibodies diluted in blocking buffer for 30 minutes (anti-rabbit Rhodamine Red-X (1:1500), anti-rabbit Cy5 (1:1500), anti-mouse AlexaFluor 488 (1:1500), anti-mouse Rhodamine Red-X (1:1500)). Slides were then washed 3 times with PBS, 0.1% Triton X-100 and mounted in mounting medium (PBS, 90% (by volume) glycerol, and 0.1 M propyl gallate). Deconvolution microscopy was performed on a DeltaVision core system (Applied Precision) equipped with an Olympus IX71 microscope, a 100x objective (NA 1.4), and a cooled charge-coupled device camera (CoolSnap HQ2; Photometrics). Images were acquired with softWoRx software (Applied Precision).

### Data collection, analysis, and statistics

Centriole counts and KD sphere measurements were performed manually on an Olympus IX71 microscope. Densitometry was performed in Fiji (ImageJ, NIH). All data were collated in Microsoft Excel and statistical analyses were performed with Prism (GraphPad). Details of statistical tests and significance information are presented in figure legends. Statistical significance was inferred when P≤0.5. Figures were prepared using Adobe Photoshop and Adobe Illustrator.

## Acknowledgements

We thank Dr. Paul Krieg for critical review of the manuscript as well as our funding sources.

## Funding

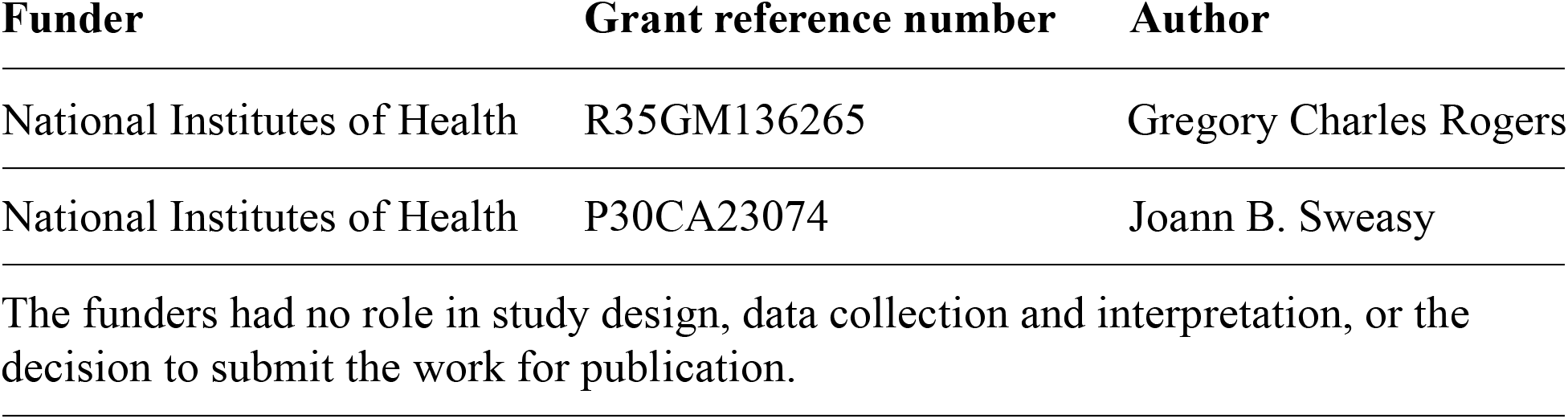

## Author Contributions

JMR was the lead in designing, performing and analyzing all experiments involving cultured fly cells as well as manuscript preparation. CJB designed and performed the experiment mapping the Asl-Plk4 interaction. AA performed the AlphaFold analysis. DWB performed the in vitro Plk4 kinase assays. SMD performed SEC-MALS analysis on purified Plk4 PB1-PB2 protein. LKS and KCS performed the crystallography and analysis of Plk4 PB3 as well as the SEC-MALS analysis of DmPB3 and HsPB3. GCR contributed to project design and manuscript preparation.

## Figure Supplement Legends

**Figure 1 – table supplement 1.** Data processing and refinement statistics for the crystal structure of *Drosophila* Polo Box 3.

| **Crystal** | **Dm Plk4 PB3 V692M SeMet** | **Dm Plk4 PB3** |
| --- | --- | --- |
| **Data Collection** |  |  |
| Wavelength (Å) | 0.97926 | 1.00000 |
| Space group | P2<sub>1</sub> | P2<sub>1</sub> |
| Unit Cell: a, b, c (Å) | 35.3, 52.1, 41.1 | 36.7, 53.1, 41.6 |
| β (°) | 106.2 | 107.9 |
| Resolution (Å) | 50.00-1.90 (1.97-1.90) | 50.00-1.75 (1.81-1.75) |
| # Reflections: Unique | 21,024 (1554) | 14,468 (961) |
| Completeness (%) | 94.8 (69.7) | 93.4 (62.4) |
| Mean redundancy | 3.4 (2.3) | 3.5 (2.4) |
| <I/σI> | 18.8 (3.9) | 22.5 (4.0) |
| R<sub>sym</sub> | 0.062 (0.270) | 0.048 (0.251) |
| **Refinement** |  |  |
| Resolution (Å) |  | 39.6-1.75 (1.81-1.75) |
| R/ R<sub>free</sub> (%) |  | 19.2 (21.9) / 23.3 (27.6) |
| # Reflections, R/R<sub>free</sub> |  | 12,766 (1420) / 746 (80) |
| Total atoms: Protein / Water |  | 1247 / 118 |
| Stereochemical ideality (rmsd):<br>Bonds (Å) / Angles (°) |  | 0.013 / 1.644 |
| Ramachandran Analysis:<br>Favored / Allowed (%) |  | 98.1 / 1.9 |
| **PDB accession code** |  | 7RL3 |

Values in parentheses indicate statistics for the highest-resolution shell.

**Figure 1 – figure supplement 1.**
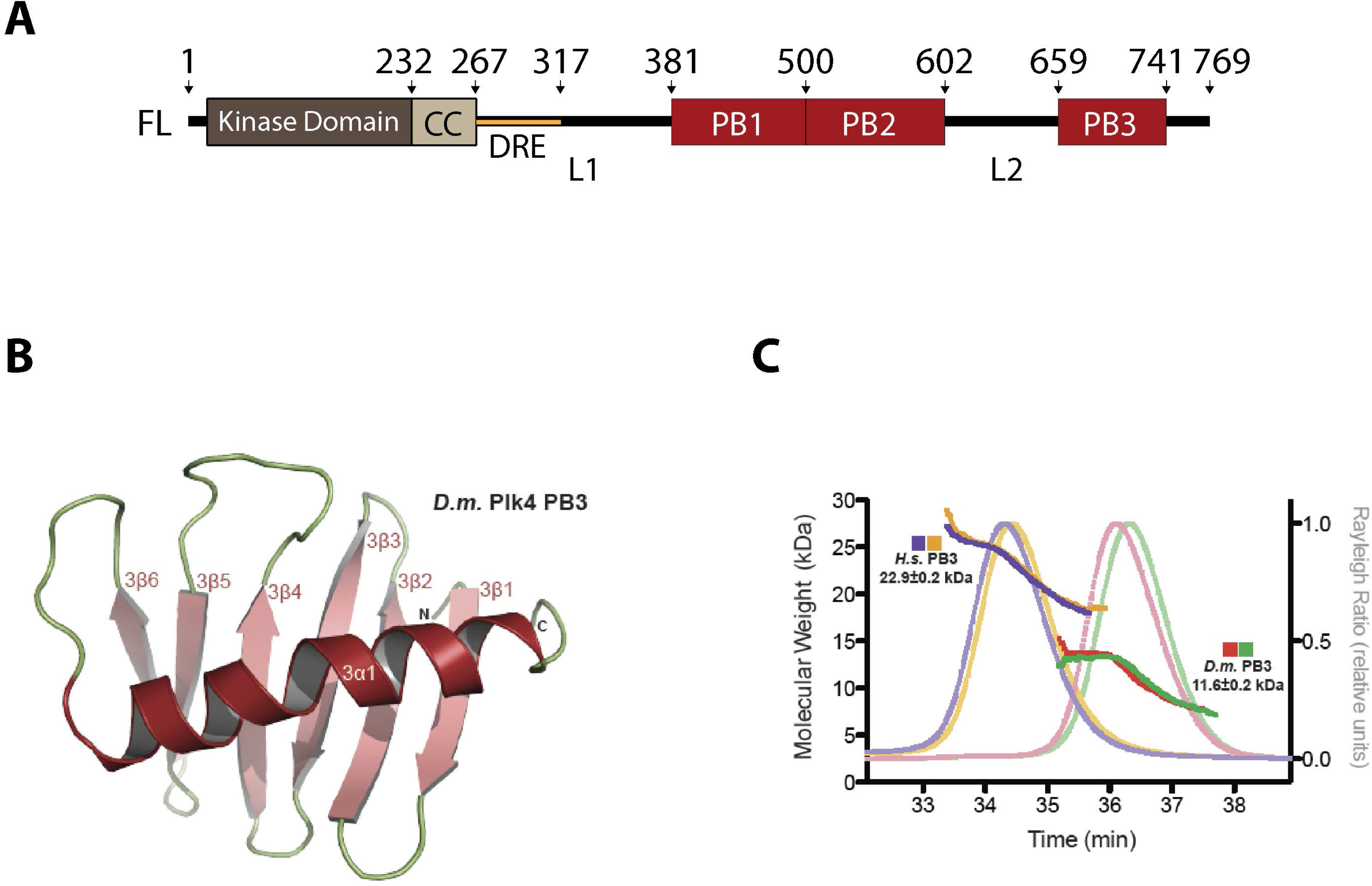
PB3 of *Drosophila* Plk4 is monomeric. (**A**) Linear map of *Drosophila melanogaster* (Dm) Plk4 depicting the functional domains. CC, coiled coil; DRE, downstream regulatory element which is a highly autophosphorylated region containing the Slimb-binding degron; PB, Polo Box; linker regions (L1 and L2). (**B**) Ribbon diagram of DmPlk4 PB3 V692M structure displaying the monomeric PB domain architecture. (**C**) *Drosophila* PB3 is not a homodimerization domain. SEC-MALS analysis of *Homo sapiens* (Hs) Plk4 PB3 and DmPlk4 PB3. The molecular mass for each construct is displayed as the average ± standard deviation. The expected monomeric size of both constructs is 11.9 kDa. Gel filtration runs were conducted at two protein concentrations: HsPB3 (blue trace: 8 mg/mL; yellow trace: 4.6 mg/mL) and DmPB3 (green trace:10 mg/mL; red trace: 27 mg/mL). HsPB3 elutes as a dimer (22.9 ± 0.2 kDa), while DmPB3 elutes as a monomer (11.6 ± 0.2 kDa).

**Figure 1 – figure supplement 2.**
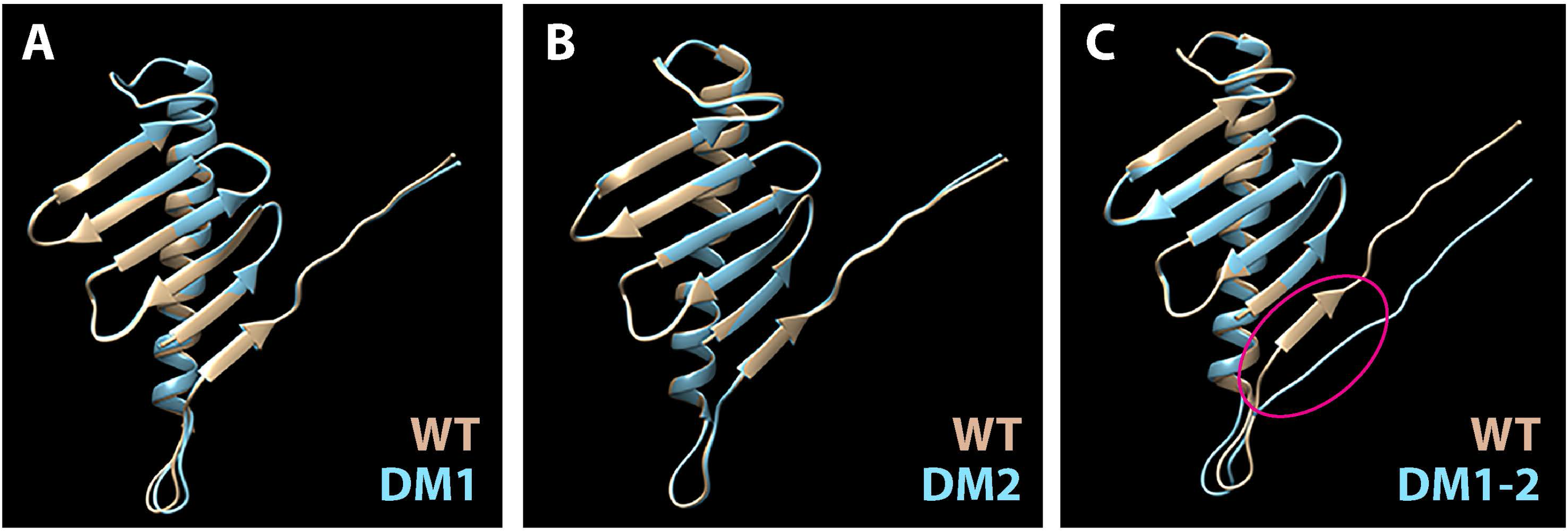
Constructs of Plk4 PB2 (aa 501-602) that harbor either dimer mutation DM1 or DM2 are predicted to adopt a native PB fold. (**A-C**) Superposition of Plk4 PB2 WT (tan) and dimer mutant (DM) (blue) predicted structures. Tertiary structures were predicted using AlphaFold and displayed using Chimera 1. Note that the structures of PB2 mutants DM1 and DM2 are nearly identical with WT structure. However, the PB2 structure of Plk4 containing all 6 substitutions (DM1-2) deviates from WT structure near its C-terminus: DM1-2 fails to form the 2β6 strand (red oval).

**Figure 2 – figure supplement 1.**
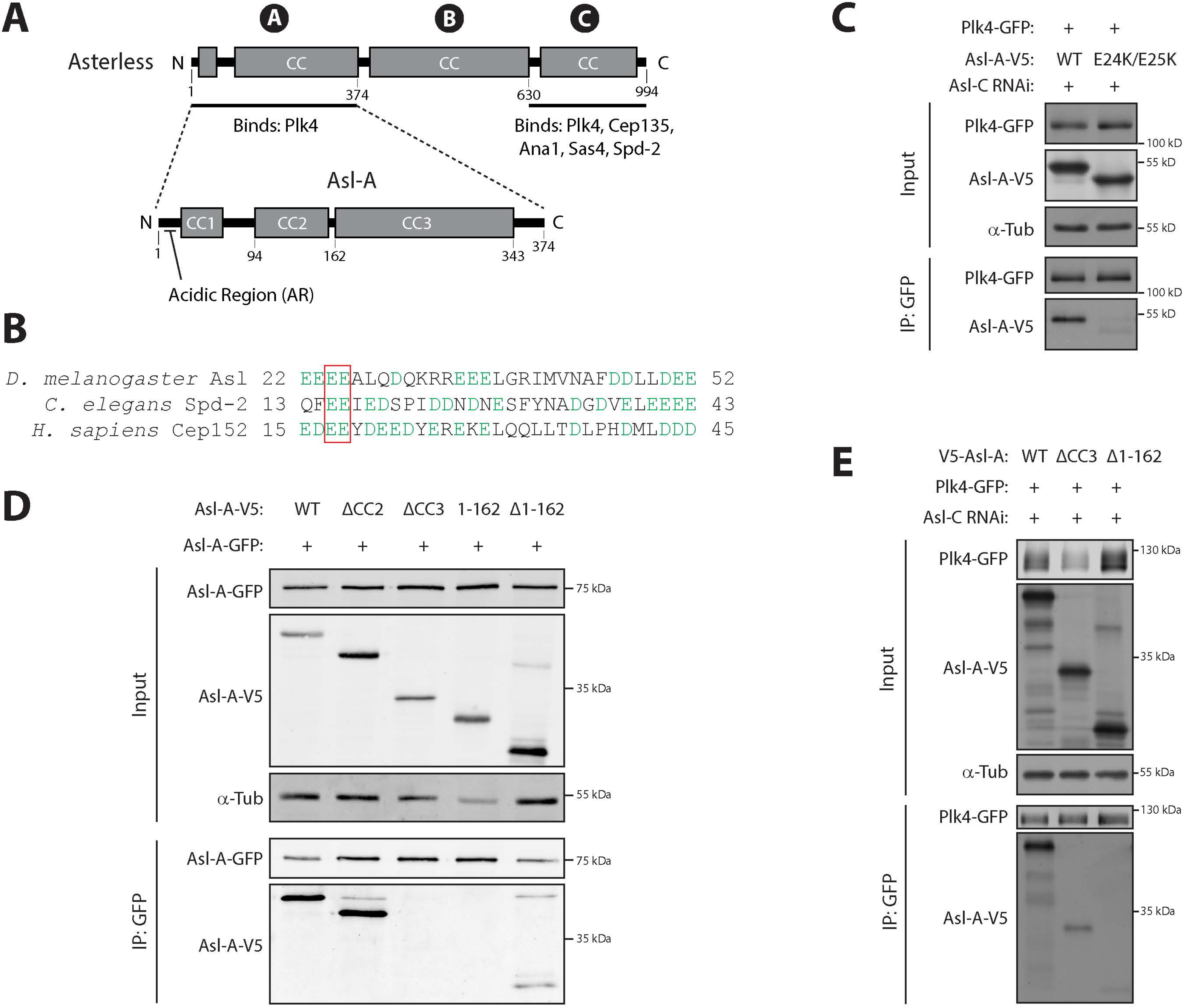
The N-terminal third of Asl (called Asl-A) contains a conserved Plk4 binding acidic region in its N-terminus and a homodimerization region in the C-terminal half of the protein. (**A**) Linear maps of *Drosophila* Asterless (Asl) showing functional and structural domains. CC, coiled-coil. The three regions of Asl are indicated with circled letters. Both Asl-A and Asl-C bind Plk4. (**B**) The N-terminus of Asl-A contains a conserved acidic region (∼30 amino acid) that binds along one face of the PB1-PB2 Z-dimer in Plk4 (Park et al., 2014; Shimanovskaya et al., 2014). Lineup of the acidic regions in the Plk4-targeting proteins Asl, Cep152 (human homolog of Asl), and *C. elegans* Spd2. Acidic residues are shown in green font. Conserved glutamic acid residues E24/E25 (red box) were mutated to lysine to generate a Plk4-binding mutant. (**C**) Mutation of the acidic region in Asl-A prevents Plk4 binding. S2 cells were Asl-depleted using RNAi targeting an exon in the gene region encoding Asl-C for 7 days. On day 4, cells were transfected with the indicated constructs and, on day 5, were induced to express for 48 hours. Anti-GFP IPs were prepared from lysates and Western blots of the samples were probed for α-tubulin, GFP, and V5. Whereas WT Asl-A co-IPs with Plk4, charge switching amino acids E24 and E25 to lysines prevents Plk4 binding. (**D**) Anti-GFP IPs were prepared from lysates of S2 cells transiently co-overexpressing the indicated inducible GFP-and V5-tagged Asl-A constructs. Blots of the input lysates and IPs were probed for α-tubulin, GFP, and V5. Note that deletion of CC3 prevents Asl-A oligomerization. (**E**) An Asl-A dimer mutant can still bind Plk4. S2 cells were Asl-depleted by RNAi targeting an exon within Asl-C for 7 days. On day 4, cells were transfected with the indicated constructs and, on day 5, were induced to express for 48 hours. Anti-GFP IPs were prepared from lysates and Western blots of the samples were probed for α-tubulin, GFP, and V5. Whereas an Asl-A mutant lacking the dimerization domain CC3 can bind Plk4, a mutant lacking the acidic region (Δ1-162) cannot co-IP with Plk4.

**Figure 2 – figure supplement 2.**
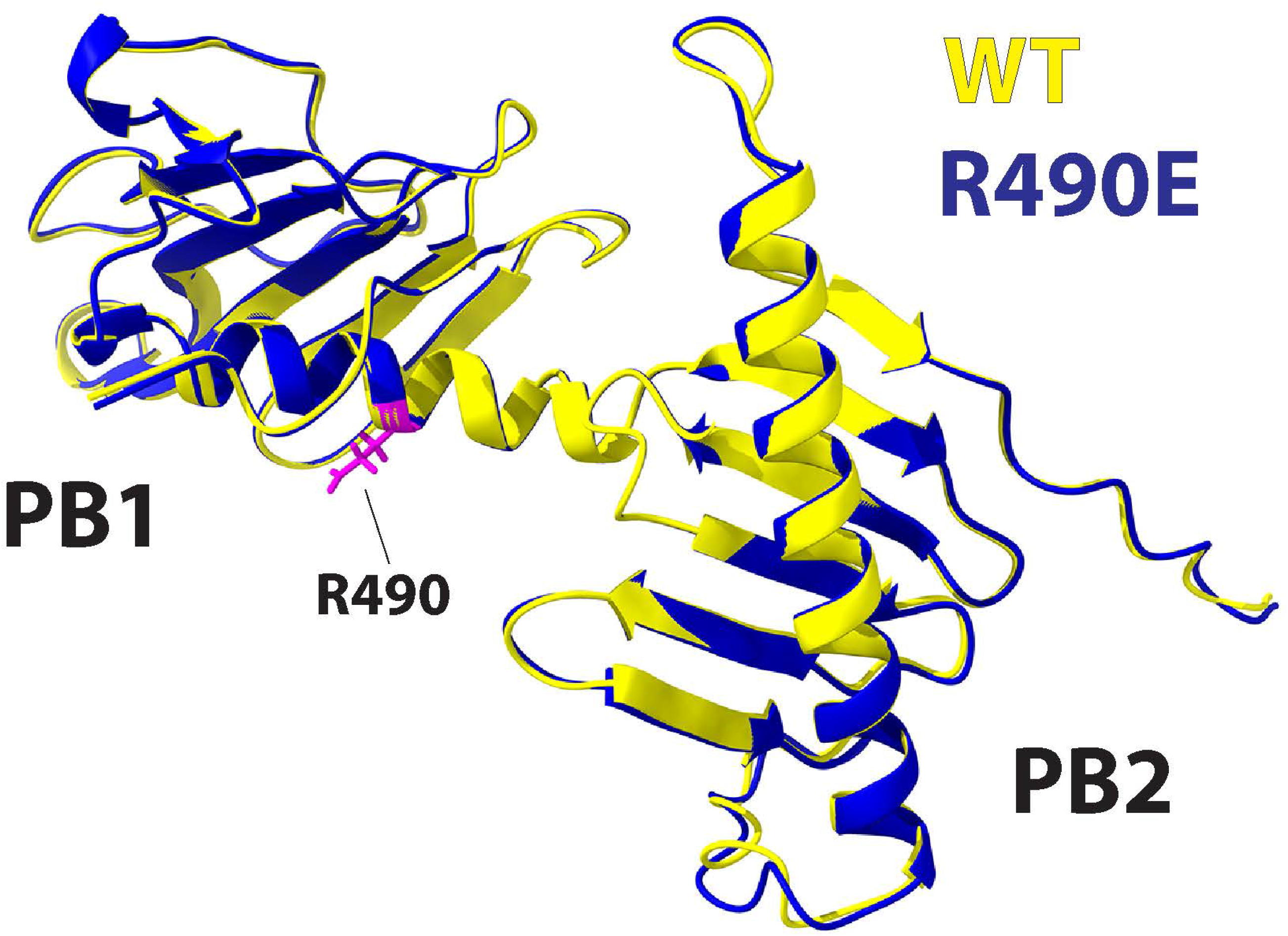
The Plk4 PB1 construct harboring the R490E substitution is predicted to adopt a native PB1-PB2 conformation. Superposition of the Plk4 PB2-PB2 structures of WT (yellow) and Asl-binding mutant R490E (blue). Tertiary structures were predicted using AlphaFold and displayed using Chimera 1. The R490 residue with side chain is shown in pink.

**Figure 3 – figure supplement 1.**
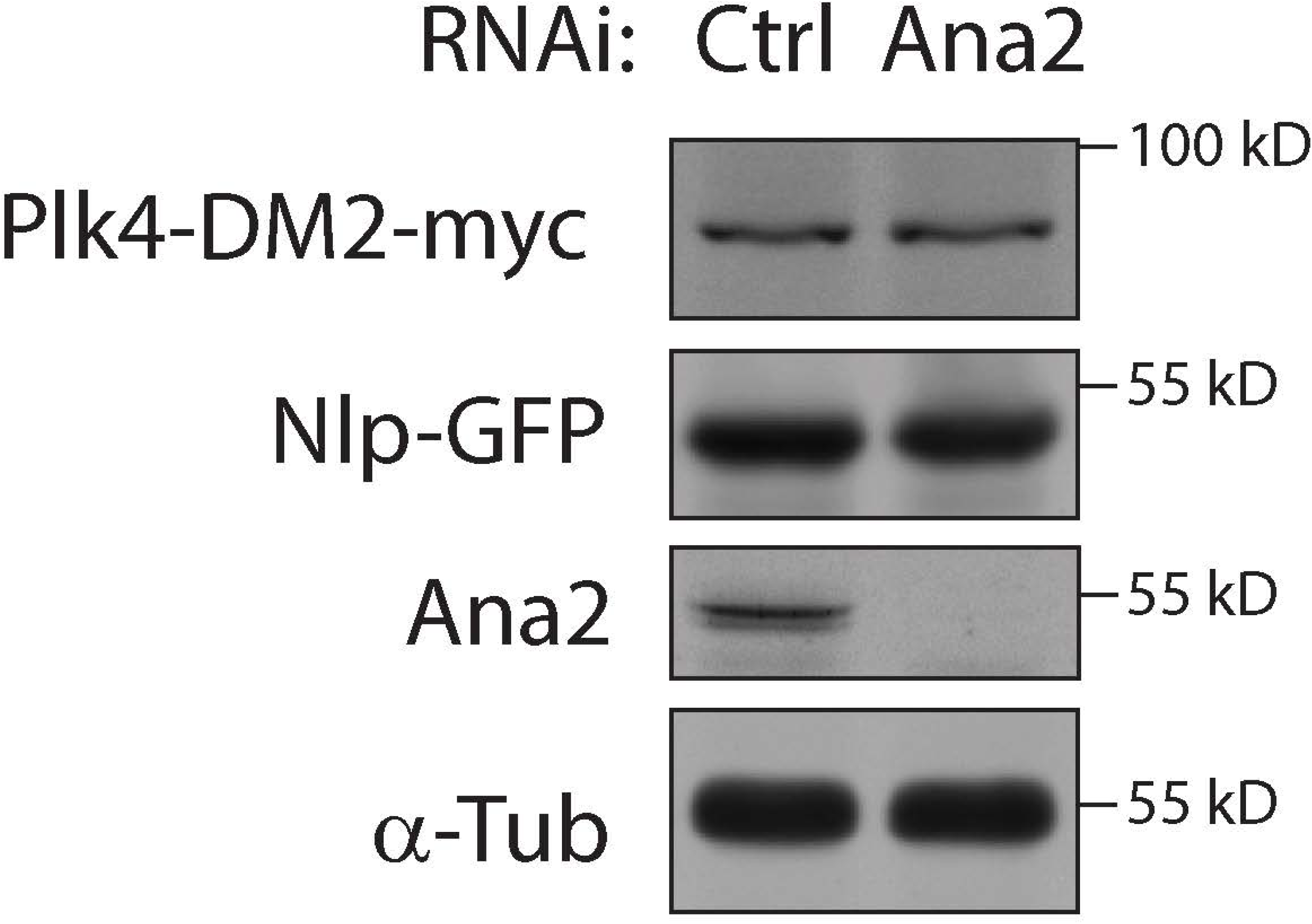
Ana2 depletion has no effect on the protein levels of dimer mutant Plk4-DM2. S2 cells were RNAi treated for 7 days. On day 4, cells were co-transfected with Nlp-GFP (Nlp: a nuclear protein used as transfection loading control) and inducible Plk4-DM2-myc. On day 5, 0.25 mM CuSO_4_ was added to the media to induce Plk4 expression. Cell lysates were prepared on day 7 and probed on immunoblots for GFP, myc, Ana2, and α-tubulin (loading control).

